# Evolution of semi-Kantian preferences in two-player assortative interactions with complete and incomplete information and plasticity

**DOI:** 10.1101/2023.05.14.540699

**Authors:** Ingela Alger, Laurent Lehmann

## Abstract

We model the evolution of preferences guiding behavior in pairwise interactions in group-structured populations. The model uses long-term evolution theory to examine different interaction scenarios, including conditional preference expression upon recognition of the partner’s type. We apply the model to the evolution of semi-Kantian preferences at the fitness level, which combine self-interest and a Kantian interest evaluating own behavior in terms of consequences for own fitness if the partner also adopted this behavior. We seek the convergence stable and uninvadable value of the Kantian coefficient, i.e., the weight attached to the Kantian interest, a quantitative trait varying between zero and one. We consider three scenarios: (a) incomplete information; (b) complete information and incomplete plasticity; and (c) complete information and complete plasticity, where individuals not only recognize the type of their interaction partner (complete information), but also conditionally express the Kantian coefficient upon it (complete plasticity). For (a), the Kantian coefficient generally evolves to equal the coefficient of neutral relatedness between interacting individuals; for (b), it evolves to a value that depends on demographic and interaction assumptions, while for (c) there are generally multiple uninvadable types, including the type whereby an individual is a pure Kantian when interacting with individuals of the same type and applies the Kantian coefficient that is uninvadable under complete information with zero relatedness when interacting with a different typed individual. Overall, our model connects several concepts for analysing the evolution of behavior rules for strategic interactions that have been emphasized in different and sometimes isolated literatures.

## 1 Introduction

This paper is about formalizing natural selection on rules guiding individual behavior in strategic interactions, a central question in evolutionary game theory (Maynard Smith and Price, 1973; Dawkins, 1980; Maynard Smith, 1982). By behavior we mean a “strategy”, i.e., “a specification of what an individual will do in any situation in which it may find itself” (Maynard Smith, 1982). In the original evolutionary game theory models, each individual is programmed to play a certain strategy regardless of the strategies used by others in the population. One way to think about this is that the strategy is innate, thus a genetically determined trait. This view led to a vast theoretical literature analysing the genetic evolution of strategies under all sorts of biological scenarios as is illustrated by the literatures on the evolution of fighting, cooperation, and life-histories in plants and animals (e.g., the books by Maynard Smith, 1982; Bulmer, 1994; Giraldeau and Caraco, 2000; Vincent and Brown, 2005; McNamara and Leimar, 2020). Here, it is the population genetic process alone that determines the “evolutionarily stable strategy” since strategies are inherited from parent to offspring and selected among alternatives by way of differential survival and reproduction.

The view that strategies are innate is restrictive, however, as it rules out situations where individuals have capacities to change their own strategy when interacting with their environment. Such processes have been incorporated into evolutionary game theory through several alternative notions, such as the concepts of “culturally stable” and “developmentally stable” strategies (Dawkins, 1980; Maynard Smith, 1982). Here, the behavior of an individual is the outcome of some updating rule(s), typically imitative or experiential, for strategy selection during the individual’s lifespan. In the memorable example detailed by Dawkins (1980), pigs in skinner boxes equilibrate on developmentally stable strategies by action reinforcement in producer-scrounger games, and a large literature has evaluated culturally stable strategies under different sorts of transmission rules (e.g., Cavalli-Sforza and Feldman, 1981; Boyd and Richerson, 1985). This in turn raises the question of what should be the evolutionarily stable rule for individual strategy selection in strategic interactions? While this question was raised early in the history of evolutionary game theory (Harley, 1981; Maynard Smith, 1982), perhaps more controversy than conclusions where initially reached (e.g., Selten and Hammerstein, 1984), and it is only more recently that this question has gained some renewed theoretical attention in evolutionary biology (e.g., Arbilly et al., 2010; Dridi and Lehmann, 2015; Dridi and Akçay, 2018; McNamara and Leimar, 2020).

In the meantime, however, economists and mathematical game theorists also produced insights about how various individual choice and transmission rules induce change in strategies in populations (e.g., the books by Sugden, 1986; Weibull, 1997; Fudenberg and Levine, 1998; Hofbauer and Sigmund, 1998; Samuelson, 1998; Young, 1998; Sandholm, 2011). One obstinate result in this literature is that updating rules of strategies–whether imitative or experiential– relying on payoff tend to converge to Nash equilibria (Hofbauer and Sigmund, 1998; Fudenberg and Levine, 1998; Cressman and Tao, 2004). Hence, in behavioral equilibrium, at the culturally stable or developmentally stable state, it is as if individuals strive to maximize the payoff function at hand and thus as if they are rational decision makers, in the sense that among a set of options they choose the one they prefer, given the others’ strategies (Mas-Colell et al., 1995). The question of what should be the evolutionarily stable rule for individual strategy selection can thus here be phrased as: if the evolving trait is the payoff function to be maximized, which payoff function is evolutionarily stable? This is the question that the literature on preference evolution for strategic interactions addresses addresses (e.g., Güth, 1995; Ok and Vega-Redondo, 2001; Dekel et al., 2007; Heifetz et al., 2007b,a; Akçay and Van Cleve, 2009; Alger and Weibull, 2010; Akçay and Van Cleve, 2012; Alger and Weibull, 2012, 2013; Wang and Wu, 2023). Because information plays a central role in strategic interactions (Fudenberg and Tirole, 1991), the formalizations of preference evolution have covered a variety of informational scenarios (e.g., Ok and Vega-Redondo, 2001; Dekel et al., 2007; Wang and Wu, 2023; see Alger and Weibull, 2019; Alger, 2023 for surveys). Focusing on the evolution of preferences gives hope to improve predictions about equilibrium behavior because payoff-based choice rules can otherwise come in endless mechanistic forms – some more biologically and cognitively inspired than others (Sutton and Barto, 1998; Russell and Norvig, 2016) but none empirically fully elucidated (e.g., Jeong et al., 2022).

The goal of this paper is to contribute to the literature on the evolution of rules guiding individual behavior in two ways and is thus divided in two parts. In the first part, we connect a number of concepts and results to analyze the long-term evolution (*sensu* Eshel, 1996; Eshel et al., 1998) of behavioral mechanisms for equilibrium action in group-structured populations. This part can thus be read as a methodological review. In the second part, we push forward within this framework the evolutionary analysis of the class of preferences involving a mix between self-interest and an interest in evaluating own behavior in the light of the consequences for own payoff if others adopted this behavior. This is the class of semi-Kantian preferences, which, in the words of Binmore (1998, p. 191), can be seen as hybrid preferences combining the categorical imperative of Nash with that of Kant. Bergstrom (1995) shows that the evolutionarily stable strategy in interactions between siblings could be interpreted as if individuals had such preferences, an interpretation that should hold more generally when interactions occur between related individuals. Semi-Kantian preferences have then indeed been shown to be evolutionarily stable and uninvadable under various transmission rules when population structure results from limited genetic or cultural mixing among interacting individuals, when interacting individuals cannot observe each other’s preferences (Alger and Weibull, 2013, 2016; Alger et al., 2020). However, so far the evolutionary convergence towards semi-Kantian preferences has not been ascertained, and their evolution has not been analyzed under different informational assumptions. Our goal is to analyse convergence stability and uninvadability of semi-Kantian preferences in three different informational scenarios: (a) incomplete information; (b) complete information and incomplete plasticity (interacting individuals can observe each other’s preferences, but an individual’s preferences do not depend on the other’s preferences); and (c) complete information and complete plasticity (interacting individuals can observe each other’s preferences, and an individual’s preferences can depend on the other’s preferences). It will be seen that the different informational and plasticity assumptions lead to quite different evolutionary outcomes, and that we are not always able to reach general conclusions about convergence stability.

Our aim is not to obtain the most general conclusions about the open questions we address, but rather to illustrate how demographic and informational features jointly contribute to the understanding of the long-term evolution of preferences in structured populations. As such, we consider only pairwise interactions and restrict attention to the parametric class of semi-Kantian preferences and the evolution of the Kantian coefficient, a quantitative trait varying between zero and one, which represents the weight attached to the Kantian interest.

## 2 Evolutionary invasion analysis of behavioral mechanisms

### 2.1 Biological assumptions for pairwise interactions

We consider a population of asexually reproducing individuals that are demographically homogeneous (no effective age, stage or sex structure). The population occupies a habitat with an infinite and constant number of groups (or demes, or spatial subdivisions), each of which is occupied by exactly two individuals and so the population is of constant size. Each individual is characterized by a type belonging to a type space Θ that affects its phenotype— the collection of any relevant morphological, physiological or behavioral measurable feature of the individual. We consider a demographic process where the population is censused at discrete time steps, between which the following events occur in cyclic order. (a) In each group, the pair of individuals engage in an interaction. Some process (learning, exchange of information, etc) leads to a pair of equilibrium strategies being expressed. The equilibrium strategy pair, which may depend on the individuals’ types as well as the types present in the population at large, determines some outcome (for example, the material payoff of each individual). (b) Each individual in each group produces a large number of juveniles according to the outcome of the pairwise interaction and eventually dies subject to some death process, ^1^ which may also depend on the outcome of the pairwise interaction. (c) Juveniles remain in the natal group with some fixed probability. With complementary probability, assumed to be non-zero, they migrate out of their natal group and survive dispersal with a certain probability that may depend on the outcome of the interaction between the juvenile’s parent and its neighbor. (d) In each group, the open reproductive spots vacated by deceased adults are randomly filled up by competing juveniles, who then become adults.

### 2.2 Invasion and individual fitness

We adopt a standard invasion analysis framework and consider a population that is monomorphic for some resident type *θ* ∈ Θ in which a mutant type *τ* ∈ Θ arises (e.g., Fisher, 1930; Eshel and Feldman, 1984; Parker and Maynard Smith, 1990; Metz et al., 1992; Charlesworth, 1994; Ferrière and Gatto, 1995; Avila and Mullon, 2023; Van Cleve, 2023). It then follows from applications of invasion analysis to our demographic process assumptions of section 2.1 (see Box 1) that any mutation *τ* ∈ Θ, which is introduced in a single individual in a monomorphic population with the resident type *θ* ∈ Θ, eventually goes extinct with probability one if and only if the invasion fitness (the geometric growth ratio) of the mutant type, denoted *W* (*τ, θ*), satisfies

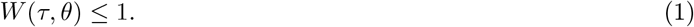

Here, the “1” can be interpreted as the growth ratio of a resident type in a monomorphic resident population, which, owing to the fact that the population is of constant size can, on average, only replace itself (i.e., *W* (*θ, θ*) = 1 for all *θ* ∈ Θ).

Invasion fitness can be represented as the individual fitness of a randomly sampled mutant *τ* descending from the individual in which the mutation initially appeared, averaged over the cases where the mutant interacts with another member of the same lineage and those where it interacts with an individual from a different lineage (who is thus of the resident type *θ*):

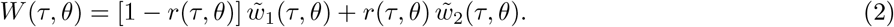

Here, 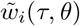 is the individual fitness of a mutant when there are *i* ∈ {1, 2} mutants in its group and the population is otherwise monomorphic for *θ*, and *r*(*τ, θ*) is the *pairwise relatedness* between a *τ* mutant and its group neighbor (see Box 1 for a derivation of eq. (2)). Pairwise relatedness is the probability that, conditional on an individual being of type *τ*, the group neighbor belongs to the same ancestral lineage and is thus also of type *τ*, whereby both individuals are *identical-by-descent* (Malécot, 1969); note that since migration is assumed non-zero, we have *r*(*τ, θ*) <1. Whether relatedness *r*(*τ, θ*) depends on both the mutant and the resident type, only on the resident type, or neither, depends on demographic and interaction assumptions. For instance, relatedness is independent of the types for family-structured populations, in which case it is determined only by the pedigree relatedness, e.g. *r* = 1*/*2 for full-siblings [as implied by the model of Michod, 1980, which also entails that eq. (2) applies to sexual reproduction in family-structured populations in the absence of inbreeding].

When *W* (*τ, θ*) is differentiable (which is not always the case), a resident type *θ*\* is locally convergence stable if and only if the first two following conditions hold, while it is locally uninvadable if the first and the third conditions hold (Eshel, 1983; Taylor, 1989; Christiansen, 1991; Geritz et al., 1998):

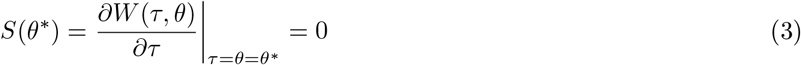

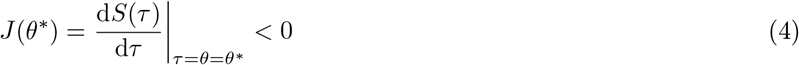

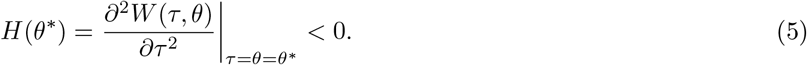

Here, *S*(*θ*), *J* (*θ*), and *H*(*θ*), stand respectively for the selection gradient, the selection Jacobian, and the selection Hessian, evaluated at the resident type *θ*. A type satisfying *S*(*θ*\*) = 0 will be called a singular type (or a singularity).^2^ There is a non-trivial relationship between the static conditions (3)-(5) obtained from invasion fitness and dynamic stability. Namely, for mutants with small effects on the phenotype, i.e. the difference |*θ* − *τ* | is small, a singular type *θ*\* satisfying conditions (4)-(5) is a (i) local attractor of the evolutionary dynamics under gradual evolution and (ii) resistant to invasion by small deviations. ^3^

### 2.3 Behavioral equilibrium

In applications of evolutionary game theory, an individual’s type is often taken to be its strategy to be applied in the interaction at hand. Yet many applications require to decouple types from strategies. In order to do this and obtain a full description of how individual fitness depends on own type and neighbor’s type — a dependence that in eq. (2) was captured through the mappings 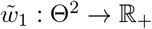 and 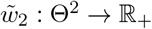, without reference to the strategies used by the individuals — we begin by defining individual fitness as a function of the strategies used, and then we introduce notation and assumptions for how the equilibrium strategies depend on the types.

Letting *χ* denote the set of strategies that each individual has access to when interacting with its neighbor, the individual fitness function *w* : *χ*^3^ → ℝ_+_ is defined such that *w*(*x*_*i*_, *x*_*j*_, *y*) gives the expected number of descendants (including the surviving self) produced over one demographic time period by an adult individual *i* expressing strategy *x*_*i*_ when matched to a group neighbour *j* expressing strategy *x*_*j*_, when individuals in the population at large all use strategy *y*. Note that any individual fitness function is subject to the demographic consistency relation *w*(*y, y, y*) = 1 for all *y* ∈ *χ* and an example thereof is provided in Box 2.

Turning now to the equilibrium strategies, in a population with a mutant type *τ* and a resident type *θ* ≠*τ*, each group either has zero, one, or two mutants. For groups with two residents (resp. two mutants), we denote by 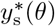 (resp. 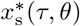) an equilibrium strategy for each individual, where the subscript “s” refers to *same* type (note that we rule out equilibria in which two identical individuals use different strategies). For mixed groups, with one resident and one mutant, let 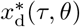 denote the mutant’s equilibrium strategy and 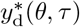 the resident’s equilibrium strategy, where the subscript “d” stands for *different* types. Importantly, throughout we assume that for any type pair (*θ, τ*) ∈ Θ^2^ with *θ* ≠ *τ*, there exist unique equilibrium strategies 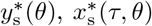, and 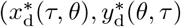). In a population with a mutant *τ* = *θ* interactions can occur between same and different lineage members having the same type. For this case, we assume that all individuals in all pairs use the same strategy 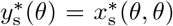 since they have the same type. This implies that the mappings 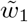 and 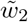 used in eq. (2) are well defined ^4^, as follows:

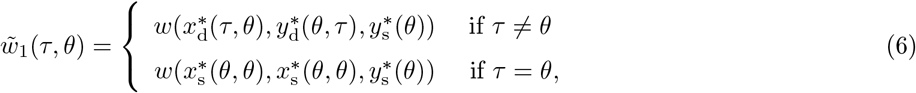

And

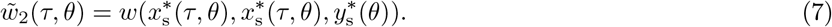

How do the equilibrium strategies arise? In the evolutionary game theory literature, a variety of processes, or mechanisms, of interdependent strategy expression have been examined, including reactive strategies, behavior response rules, learning rules, or developmental rules (e.g., Maynard Smith, 1982; McNamara et al., 1999; Akçay and Van Cleve, 2009; Killingback and Doebeli, 2002; Taylor and Day, 2004; André and Day, 2007; Dridi and Akçay, 2018; McNamara and Leimar, 2020). In each case, a dynamical system drives strategy expression over time, and under some conditions these behavioral dynamics reach an equilibrium. One way to formalize these mechanisms is to posit that the equilibrium strategies solve a fixed-point problem. Thus, for mixed groups, let there be two mappings, *M*_d_ : Θ^2^ × *χ* ^2^ → ℝ for the mutant type and *R*_d_ : Θ *× χ* ^2^ → ℝ for the resident type, which capture the process at hand, and which are such that an equilibrium pair of strategies satisfies the fixed-point system of equations:

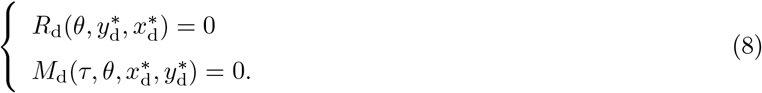

The mechanism *M*_d_ is parametrized by both the mutant and the resident type, while *R*_d_ is parametrized only by the resident type. This is so because when individuals interact their strategy may depend on (i) their own type and strategy, (ii) the strategy of their interaction partner, and (iii) on strategies in the population at large, which depends only on the resident type when the mutant is rare (see also eq. 10 below). Solving for 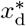 and 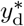 produces the dependence of each strategy on both types, i.e., 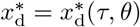 and 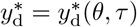.

For the equilibrium strategy used in mutant-mutant interactions, let there be a mapping *M*_s_ : Θ^2^ *× χ* → ℝ which describes the process whereby a mutant interacts with another mutant as a function of the partner’s strategy. The strategy in equilibrium is then assumed to satisfy the fixed-point equation

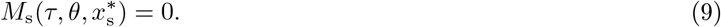

The behavioral mechanism *M*_s_ is parametrized by both the mutant and the resident type, because the strategy used in the population at large (which depends on the resident type) may affect the strategy used in a mutant-mutant pair. Hence, the solution of eq. (9) implicitly defines the equilibrium strategy as a function of both *τ* and *θ*, so that we can write 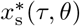. Finally, for the equilibrium strategy in resident-resident interactions, the equilibrium strategy 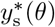 is assumed to solve the fixed-point equation

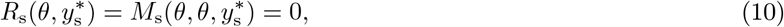

where *R*_s_ : Θ*×χ* → ℝ is the behavioral mechanism characterizing the (same) equilibrium strategy of each individual in a resident pair. By contrast to the equilibrium strategy between two mutants, which depends both on the mutant and the resident type, the equilibrium strategy between two residents depends only on the resident type.

### 2.4 Nash equilibrium and utility function

Many formalizations of the behavioral fixed points (8)–(10) consist in assuming that strategies equilibrate by being guided by some payoff function and adopting assumptions such that the dynamics lead to a Nash equilibrium according to this payoff function. In such models, the mappings *R*_s_, *R*_d_, *M*_s_, and *M*_d_ can be thought of as describing the best response functions according to the payoff function. In equilibrium, it is thus as if individuals maximize this payoff function, given the strategy used by the opponent. One class of such models takes the payoff function to be a *utility function*, which represents an individual’s preferences. ^5^ In our setting, given some resident utility function determined by *θ* and some mutant utility function determined by *τ*, the strategy 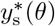 (resp. 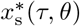) would be the strategy in *χ* that maximizes the utility of a resident (resp. that of a mutant), given that the resident (resp. mutant) with whom it interacts also uses strategy 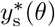 (resp. 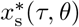). Likewise, the strategy 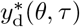 would be the strategy in *χ* that maximizes the utility of the resident, given that the mutant with whom it interacts uses strategy 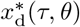, while the strategy 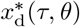) would be the strategy in *χ* maximizing the mutant’s utility, given that the resident with whom it interacts uses strategy 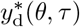.

We endorse this approach and rely on results showing that among the set of all continuous utility functions, a utility function representing semi-Kantian preferences emerges as being particularly viable from an evolutionary perspective (Alger and Weibull, 2013; Alger et al., 2020). For some individual who uses strategy *x* when it neighbour uses strategy *y*, and given that strategy *y*\* is played at large in the population, this utility function is defined as

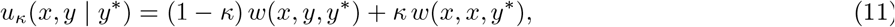

where *w* is the individual fitness function defined above and *κ* ∈ [0, 1]. The first is the individual’s realized fitness, given the strategies used. The second term is the fitness that the individual would realize if – hypothetically – the opponent used the same strategy (*x*) instead of strategy *y*; since the individual thereby evaluates what would happen if others were to follow the same course of action as itself, the second term can be interpreted as capturing a form of the first formulation of Kant’s categorical imperative: “act as if the maxims of your action were to become through your will a universal law of nature” (Kant, 1785). These preferences were dubbed *Homo moralis* (Alger and Weibull, 2013), yet they apply regardless of the organism under consideration in our life-cycle assumptions of section 2.1. We will call the parameter *κ* the Kantian coefficient and our goal is to investigate its evolution under three different scenarios: (a) incomplete information, (b) complete information with incomplete plasticity, and (c) complete information with complete plasticity. Each of these scenarios, together with the utility function (11), defines a specific set of behavioral mechanisms (8)–(10), detailed in the next section.

## 3 Evolution of the Kantian coefficient

For simplicity, we restrict attention to settings where *w* is twice continuously differentiable and the utility function (11) is strictly concave in its first argument for any *θ* ∈ [0, 1]. We further take the strategy space *χ* to be an open and convex subset of ℝ. These assumptions together imply that any equilibrium strategy must satisfy first-order conditions, and this facilitates the analysis. To rule out trivial settings in which an individual’s strategy has no impact on the opponent’s fitness, we also assume that *∂w*(*x, y, z*)*/∂y* ≠ 0 for all (*x, y, z*) ∈ *X*^3^. We further assume that the sign of this effect is independent of the strategies used, and by convention, we let *∂w*(*x, y, z*)*/∂y >* 0 for all (*x, y, z*) ∈ *X*^3^, meaning that an increase in the strategy of an individual’s partner enhances the individual’s fitness.^6^. Finally, we assume that *∂*^2^*w*(*x, y, z*)*/∂y∂x* has the same sign for all (*x, y, z*) ∈ *X*^3^, and we will say that the strategies are *strategic complements* if *∂*^2^*w*(*x, y, z*)*/∂y∂x* > 0, *strategic substitutes* if *∂*^2^*w*(*x, y, z*)*/∂y∂x* <0, and *strategically neutral* if *∂*^2^*w*(*x, y, z*)*/∂y∂x* = 0.

### 3.1 Incomplete information

#### 3.1.1 Behavioral equilibrium

Under incomplete information, an individual’s type is the value of their Kantian coefficient taking a value in the interval [0, 1], and an individual cannot observe the type of its interaction partner. Still, the individual can have information about the matching distribution in the pairwise interaction, i.e., the probability that the partner belongs to the same lineage. One-shot interactions between perfect strangers are examples of this kind of interaction, as are interactions between family members when the only available information is their degree of kinship. We assume that individuals hold the belief that the probability of being matched with an individual from the same lineage is given by *r*(*τ, θ*), which is a correct belief in the sense that a randomly drawn mutant in the lineage started by the initial mutant, faces the probability *r*(*τ, θ*) of being matched with another mutant. Given these assumptions, an individual can condition its strategy only on the strategy that it expects its partner to use, given the belief on the matching distribution. Since any individual uses the same strategy whether the neighbor has the same or a different type, we simplify the notation by setting 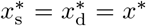 and 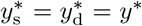, where *x*\* is the equilibrium strategy of mutants and *y*\* that of residents. A strategy pair (*x*\*, *y*\*) is a (Bayesian) Nash equilibrium if (a) *y*\* is a preferred strategy for a resident, given that other residents use strategy *y*\*; and (b) *x*\* is a preferred strategy for a mutant, given that residents use *y*\* and the other mutants use *x*\*, and given that the mutant applies the belief that the probability of being matched with another mutant is

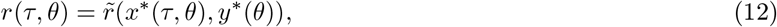

where on the right-hand side relatedness is expressed in terms of the equilibrium strategies of mutant and resident individuals. Formally 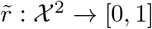 so that 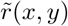 is the relatedness of a mutant towards a random group member when mutants play strategy *x* and residents play strategy *y* [a concrete example thereof is obtained by setting 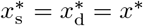 and 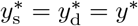 into the right-hand side of eq. (B-k) of Box 2]. Thus, (*x*\*, *y*\*) solves the fixed point system

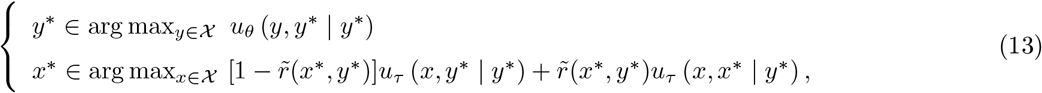

which is fully in line with the model in Alger et al. (2020, eq. 1 and eq. 5) and where the utility functions *u*_*θ*_ and *u*_*τ*_ are defined in eq. (11).^7^ The behavioral fixed point (13) defines the behavioral mechanisms (8)–(10), which here satisfy *M*_d_(*τ, θ, x*\*, *y*\*) = *M*_s_(*τ, θ, x*\*) = 0 and *R*_d_(*θ, y*\*, *x*\*) = *R*_s_(*θ, y*\*) = 0, since an individual’s strategy choice cannot be conditioned on the interactant’s type.

Under our mathematical assumptions, the (assumed unique) equilibrium pair of strategies satisfies the necessary first-order conditions for the maximization problems in eq. (13):

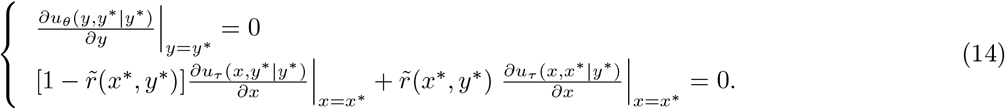

Using eq. (11), these equations become

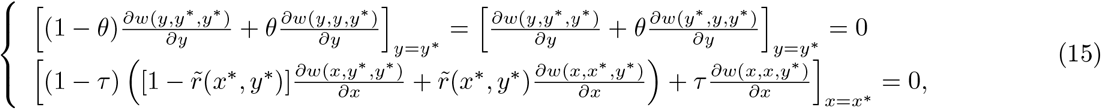

where the second equality of the first line shows that the marginal change in utility can be expressed as the sum of the effect of own behavior on own fitness and the fitness of the partner weighted by the Kantian coefficient *θ*. The necessary second-order conditions for (*x*\*(*τ, θ*), *y*\*(*θ*)) defined by (14) to be maxima rather than minima are:

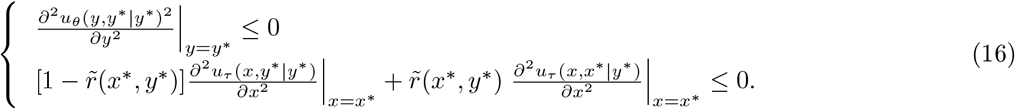

These inequalities hold strictly by virtue of the assumption that *u*_*θ*_ is strictly concave in its first argument. Hence:

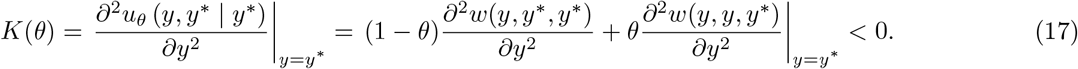

This inequality can in turn be used to evaluate how the mutant’s equilibrium strategy would change if the mutant trait value changed. To see this, by applying the implicit function theorem, one obtains by totally differentiating the second line of eq. (15) with respect to *τ* and solving the resulting linear equation for *∂x*\*(*τ, θ*)*/∂τ* :

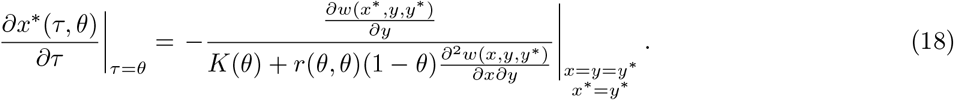

This (local) *mutant behavioral perturbation*, which will be seen to play a central role in the evolutionary analysis, is always positive, i.e., *∂x*\*(*τ, θ*)*/∂τ* |_*τ*=*θ*_ *>* 0 (see Appendix A for a proof). The intuition is that by making the individual attach a greater value to the effect that the strategy would have on self if it was also adopted by the other individual, an increase in *τ* makes the individual internalize the positive externality (*∂w*(*x, y, y*\*)(*∂y*) *>* 0) that the other’s strategy has on self.

#### 3.1.2 Evolutionary equilibrium

Turning now to the analysis of selection on the Kantian coefficient, invasion fitness writes

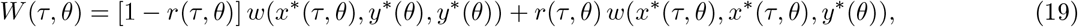

which is differentiable (since individual fitness is differentiable so will be *r*(*τ, θ*), see Box 1). Then, substituting eq. (19) into *S*(*θ*) = *∂W* (*τ, θ*)*/∂τ* |_*τ*=*θ*_, the selection gradient is

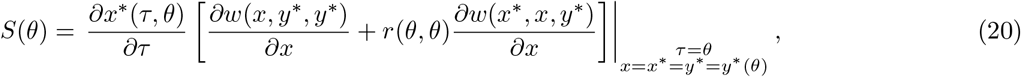

because the term multiplying *∂r*(*τ, θ*)*/∂τ* is *w*(*x*\*(*τ, θ*), *x*\*(*τ, θ*), *y*\*(*θ*))|_*τ*=*θ*_ − *w*(*x*\*(*τ, θ*), *y*\*(*θ*), *y*\*(*θ*))|_*τ*=*θ*_ = 0. Selection on the Kantian coefficient thus depends on the (local) mutant behavioral perturbation weighted by the inclusive fitness effect at the strategy level, i.e. the sum of the direct effect, *∂w*(*x, y*\*, *y*\*)*/∂x*, and the indirect effect, *∂w*(*x*\*, *x, y*\*)*/∂x*, on fitness weighted by neutral relatedness. Moreover, since *∂w*(*x, y*\*, *y*\*)*/∂x* = *−θ∂w*(*y*\*, *y, y*\*)*/∂y* at *τ* = *θ* and *x*\*= *y*\*(see eq. (15)), we obtain that

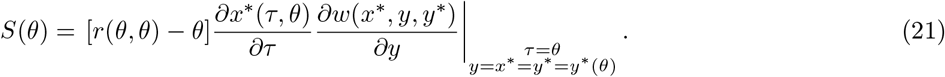

Since (by assumption) *∂w*(*x, y, y*\*)*/∂y* 0 for all (*x, y, y*\*) ∈ *χ*^3^, which also implies that *∂x*\*(*τ, θ*)*/∂τ ≠* 0 (see eq. (18)), this selection gradient shows that the unique singular trait value is

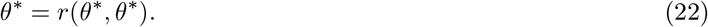

But is *θ*\*convergence stable and uninvadable?

Let us first consider convergence stability, by determining whether the Jacobian

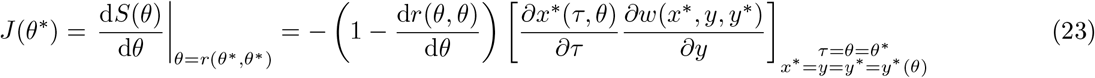

is strictly negative. Since the term in the square brackets is strictly positive, we immediately obtain that *θ*\*= *r*(*θ*\*, *θ*\*) is convergence stable if and only if d*r*(*θ, θ*)*/* d*θ* <1.

What about local uninvadability? To ascertain this, we examine whether the Hessian is strictly negative. Given that *∂w*(*x, y*\*, *y*\*)*/∂x* = *−θ∂w*(*y*\*, *x, y*\*)*/∂x* when *τ* = *θ* (as noted already above), we obtain:

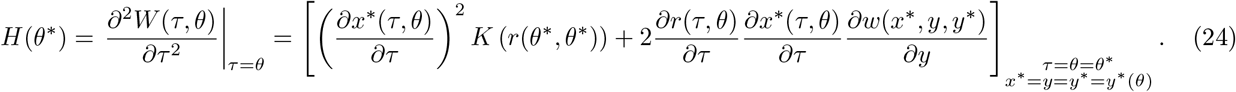

Since *K* (*r*(*θ*\*, *θ*\*)) <0, the first term is strictly negative. Hence, a sufficient condition for *θ*\*= *r*(*θ*\*, *θ*\*) to be (locally) uninvadable is that the local perturbation of relatedness, *∂r*(*τ, θ*)*/∂τ*, be nil. The relatedness perturbation can be different from zero, however (for example, see eq. (B-j) for the expression of *r*(*θ, τ*) for a Moran process), and its sign typically depends on demographic and interaction assumptions in non-trivial ways. Moreover, it does not involve second-order derivatives of individual fitness (Mullon et al., 2016), and thus does not vary systematically according to the strategic substitutability or complementarity of the strategies. Hence, in settings where behavior affects relatedness *∂r*(*τ, θ*)*/∂τ* ≠ 0, it is challenging to identify general conditions that would guarantee that *J* (*θ*\*) <0 and *H*(*θ*\*) <0. Yet, it is known that in certain settings (summarized below) both d*r*(*θ, θ*)*/* d*θ* and *∂r*(*τ, θ*)*/∂τ* are negligible. We refer to this as *weak trait effects on relatedness*. We can then summarize sufficient conditions for the Kantian coefficient to be evolutionarily stable as follows, where

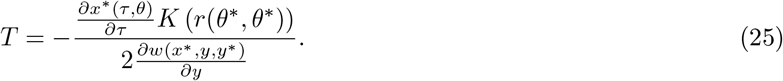

##### Result 1.

*When interactions take place under incomplete information, the Kantian coefficient equal to the neutral relatedness, θ*\*= *r*(*θ*\*, *θ*\*), *is the unique singular trait value. It is both an evolutionary attractor (convergence stable) and locally uninvadable if trait effects on relatedness are sufficiently weak. More precisely, θ*\*= *r*(*θ*\*, *θ*\*) *is convergence stable if and only if* d*r*(*θ, θ*)*/* d*θ* <1 *and it is locally uninvadable if and only if ∂r*(*τ, θ*)*/∂τ* <*T*.

While the condition for uninvadability is consistent with the results of Alger et al. (2020),^8^ our analysis reinforces those results by identifying a sufficient condition for the partly Kantian coefficient equal to neutral relatedness to be also convergence stable. When these conditions are satisfied, individuals in a population at the evolutionary equilibrium will thus behave according to Hamilton’s (marginal) rule at the strategy level, i.e., their behavioral equilibrium *y*\* satisfies

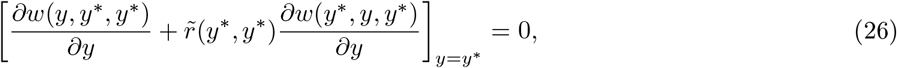

where the value of relatedness may depend endogenously on the strategy.

Interestingly, many biological scenarios do exhibit weak, or even nil, trait effects on relatedness whereby 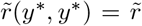 is independent of *y*\* in eq. (26).^9^ First, in family-structured populations, which cover a large class of interactions (e.g., parent-offspring interactions, interactions between sibling or cousins, etc…), relatedness is independent of the types (d*r*(*θ, θ*)*/* d*θ* = *∂r*(*τ, θ*)*/∂τ* = 0). Second, relatedness is also independent of the types in spatially-structured populations when selection is weak in the sense that the strategies in the interaction affect fitness only marginally (see, e.g., Alger et al., 2020). Such independence can extend to cases where effects are not so marginal because when the migration probability is exogenous, both d*r*(*θ, θ*)*/dθ* and *∂r*(*τ, θ*)*/∂τ* tend to be negligible for several games (Wakano and Lehmann, 2014; Mullon et al., 2016). Finally, for certain demographic processes, like the Moran process when behavior affects only reproduction, one has d*r*(*θ, θ*)*/* d*θ* = 0 and *∂r*(*τ, θ*)*/∂τ* = 0 (Mullon et al., 2016), but, as implied by eq. (B-k) of Box 2, the relatedness perturbation is non-zero in the Moran process when behavior affects survival.

### 3.2 Complete information with incomplete plasticity

#### 3.2.1 Behavioral equilibrium

Under complete information individuals have information about the type of their interaction partner, which is here again taken to be the partner’s Kantian coefficient taking values in the interval [0, 1]. However, an individual’s own utility function cannot be conditioned on that information: this is what we mean by incomplete plasticity. Because individuals can observe the type composition of their group, whenever the mutant type differs from the resident type, the distinction between a mutant’s equilibrium strategies 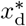 and 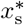, as well as between a resident’s equilibrium strategies 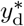 and 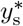, is relevant, as per the behavioral mechanisms (8)–(10). Hence, the equilibrium strategy 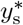 used by each resident in an interaction with another resident satisfies

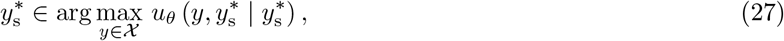

the equilibrium strategy 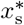 used by each mutant in an interaction with another mutant satisfies

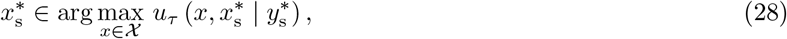

and the equilibrium pair of strategies 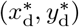 used by a mutant and a resident, respectively, in a mutant-resident interaction solves the fixed point system

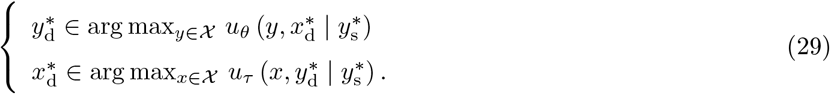

The fixed point equations (27)–(29) define the behavioral mechanisms (8)–(10) under complete information with incomplete plasticity. Note that if *τ* = *θ* in eq. (29), then 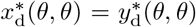 owing to the strict concavity of *u* in its first argument, which implies that to each strategy played by the opponent there exists a unique best response. Hence, if *τ* = *θ*,

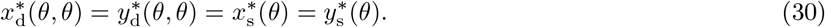

As in the incomplete information scenario, in the evolutionary analysis we use the expressions that capture how the equilibrium strategies are modified by marginal changes in the mutant trait and the resident trait. To obtain these behavioral perturbations, we first write the necessary first-order conditions for 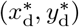 to be a Nash equilibrium:

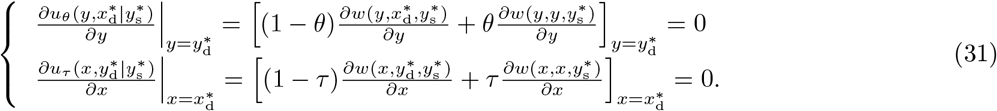

Therein, the monomorphic resident behavioral equilibrium 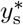 solves

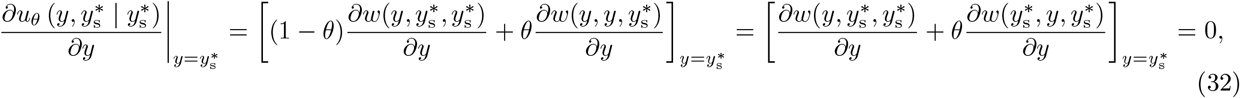

where the second equality shows that the marginal change in utility in the resident population can, as under incomplete information, be expressed as the sum of the effect of own behavior on own fitness and the fitness of the partner weighted by the Kantian coefficient *θ*. Since *u*_*θ*_ is strictly concave in its first argument, the second-order partial derivative of *u*_*θ*_, evaluated at 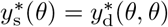, is strictly negative:

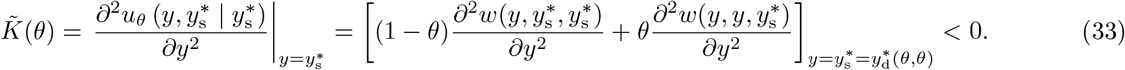

The system of equations in eq. (31) together implicitly define 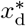 and 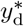 as functions of *τ* and *θ*. Applying the implicit function theorem, we obtain the following expressions for the behavioral perturbation of the equilibrium strategy of a mutant and of a resident with respect to the mutant trait value, evaluated locally at *τ* = *θ*:

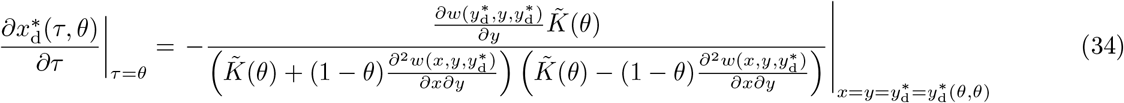

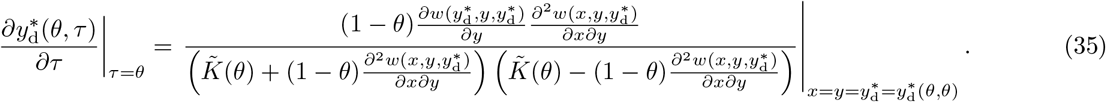

In the evolutionary analysis, it is the ratio of the resident’s to the mutant’s behavioral perturbation that will matter:

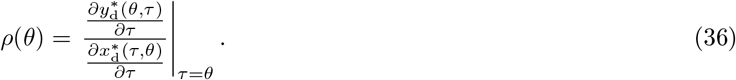

This is well defined, since the assumption 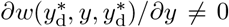 implies 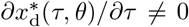, which means that the equilibrium strategy of mutants always changes as a result of a marginal change in the mutant trait value. Because *ρ*(*θ*) measures the extent to which an individual’s neighbour’s strategy varies with own strategy variation, we follow previous terminology and refer to *ρ*(*θ*) as the *response coefficient* (Akçay and Van Cleve, 2012). *Inserting eq. (34) and eq. (35) into eq. (36), we* obtain

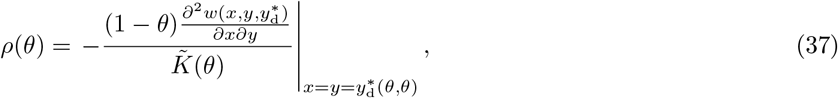

implying that the response coefficient has the same sign as 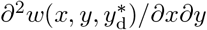, and this will play a role in the analysis of selection on the Kantian coefficient, to which we now turn.

#### 3.2.2 Evolutionary equilibrium

To begin, note that eq. (30) implies that we can write invasion fitness (2) as follows:

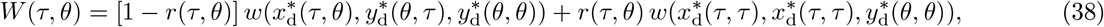

which is differentiable. Substituting this into the selection gradient *S*(*θ*) = *∂W* (*τ, θ*)*/∂τ* |_*τ*=*θ*_, and simplifying yields (since the term multiplying *∂r*(*τ, θ*)*/∂τ* is 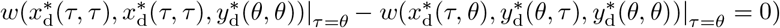

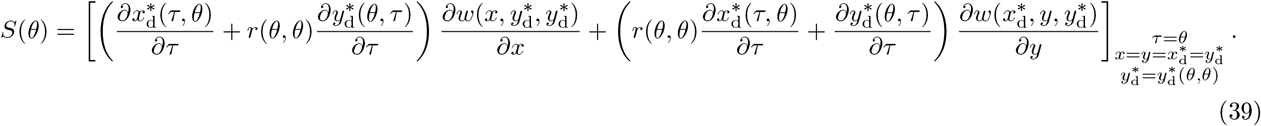

From eq. (32), we can write 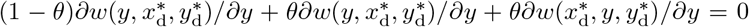 at *τ* = *θ* where 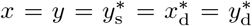. Therefore, we can replace 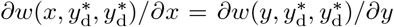 by 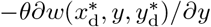 in eq. (39), to obtain

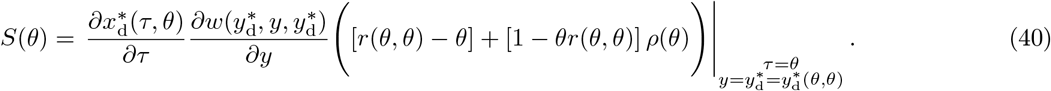

Comparing eq. (40) to the selection gradient under incomplete information (see eq. (21)), we see that if the response coefficient is nil, i.e., if *ρ*(*θ*) = 0, the two selection gradients are identical, and *θ* = *r*(*θ, θ*) is then the unique singularity. This is not surprising since under incomplete information changes in the mutant trait value has no effect on the resident’s equilibrium strategy. We further observe that when *θ* = 1, then *ρ*(*θ*) = 0; using eq. (34) in eq. (40), the selection gradient at *θ* = 1 is thus

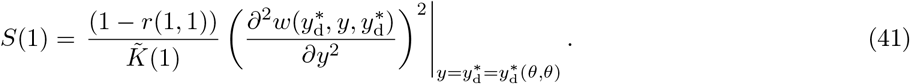

Since 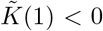 and *r*(*θ, θ*) <1 for all *θ* ∈ [0, 1], we obtain *S*(1) <0, which implies that *θ* = 1 is always counterselected, and can neither be convergence stable nor uninvadable. By contrast, nothing allows to rule out that *θ* = 0 could be convergence stable and/or uninvadable. More generally, since 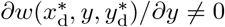 (by assumption), eq. (40) implies that *S*(*θ*) = 0 if and only if

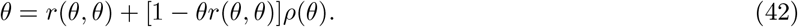

Let 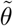 denote a solution to this equation. Since *r*(*θ, θ*) <1 for all *θ* ∈ [0, 1], so that 1 *− θr*(*θ, θ*) *>* 0 for any *θ* ∈ [0, 1], it follows immediately from eq. (42) that 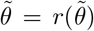 if 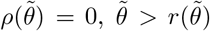 if 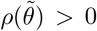, and 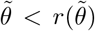. Recalling that the sign of *ρ*(*θ*) depends on the sign of 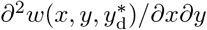 (see eq. (37)), and that we restrict the Kantian coefficient to take values between 0 and 1, the following result obtains.

##### Result 2.

*Let θ*\**denote a singularity for the Kantian coefficient under complete information and incomplete plasticity, and* 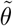 *denote a solution to eq*. (42). *Then:*

1. *θ*\*= 0 *if* 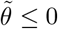, *which requires w to be such that strategies are strategic substitutes or strategically neutral;*
2. 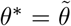 *if* 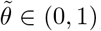, *in which case θ*\*= [*r*(*θ*\*, *θ*\*) + *ρ*(*θ*\*)]*/*[1 + *ρ*(*θ*\*)*r*(*θ*\*, *θ*\*)]. *In particular, θ*\*= *r*(*θ*\*, *θ*\*) *if w is such that strategies are strategically neutral, while θ*\*<*r*(*θ*\*, *θ*\*) *(resp. θ*\**> r*(*θ*\*, *θ*\*)*) if w is such that strategies are strategic substitutes (resp. complements), and θ*\*= *ρ*(*θ*\*) *if r*(*θ*\*, *θ*\*) = 0.

By contrast to the incomplete information setting where the Kantian coefficient must coincide with the coefficient of relatedness, here it can be either larger or smaller and the behavioral equilibrium *y*\* in a population at an evolutionary equilibrium *θ*\*now satisfies

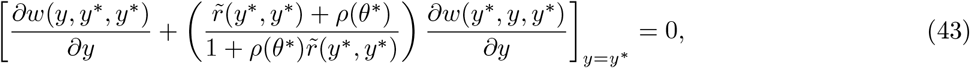

where relatedness will be independent of the strategies (and types) under weak trait effects on relatedness. Whether the Kantian coefficient exceeds or falls short of relatedness depends on whether the fitness function exhibits, respectively, strategic complementarity or substitutability. The reason that strategic complementarity raises the Kantian coefficient above the value of relatedness stems from the fact that a mutation consisting in an increase in the Kantian coefficient then induces a positive correlated response in the strategy expression by its neighbor, implying that the marginal benefit of the Kantian coefficient is larger than under strategic neutrality (compare eq. (21)–eq. (40)). By contrast, under strategic substitutability, a mutation consisting in an increase in the Kantian coefficient has a negative impact on its neighbor’s equilibrium strategy, implying that the marginal benefit marginal benefit of the Kantian coefficient is smaller than under strategic neutrality.

Result 2 further shows that a singular Kantian coefficient can in principle take any value in the range [0, 1) depending on demographic and behavioral parameters. Interestingly, eq. (42) along with eq. (36) is identical to the corresponding equation in the model of Alger and Weibull (2012) (see their eq. (29)), wherein they examine the class of other-regarding utility functions whereby an individual may attach some evolving weight *α* ∈ (−1, 1) to the interactant’s individual fitness. Hence, Theorem 1 of this previous work also establishes that whether the exact value of the evolving weight *α* exceeds or falls short of relatedness depends on whether the fitness function exhibits strategic substitutability, complementarity, or neutrality. ^10^

We now examine whether a Kantian coefficient *θ*\* of Result 2 is convergence stable and uninvadable. Due to the complexity of the expressions for the Jacobian *J* (*θ*\*) and the Hessian *H*(*θ*\*) coefficients at *θ*\* solving *S*(*θ*\*) = 0, presented in Appendix B, we were unable to reach generic answers to these questions, and further assumptions may be needed to reach more definite results. However, we verify that convergence stability and uninvadability can obtain, by resorting to an illustrating example. Consider a Moran demographic process (i.e. individual fitness takes the form of eq. (B-i)) with constant death rate *μ* and juvenile survival probability *s*, and that individual face a pairwise interaction such that their expected fecundity (number of offspring produced at stage (b) of the life cycle of section 2.1) is linear-quadratic in the two players’ actions:

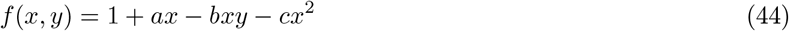

for parameters *a, b, c* ∈ ℝ. Then, substituting eq. (44) into individual fitness (B-i), we can evaluate the selection gradient (40), the Jacobian (A-10) and the Hessian (A-12) coefficients. Even for this simple example, eq. (42) is a quartic function that cannot be solved explicitly and so we analyse the selection gradient numerically. Fig. (1) displays how for fixed but different values of the backward migration rate *m*_b_ (eq. B-l), *θ*\* varies when *b* is varied. Fig. (1) shows that by depending on *b*, the Kantian coefficient takes a value above or below that of relatedness.

**Figure 1:**
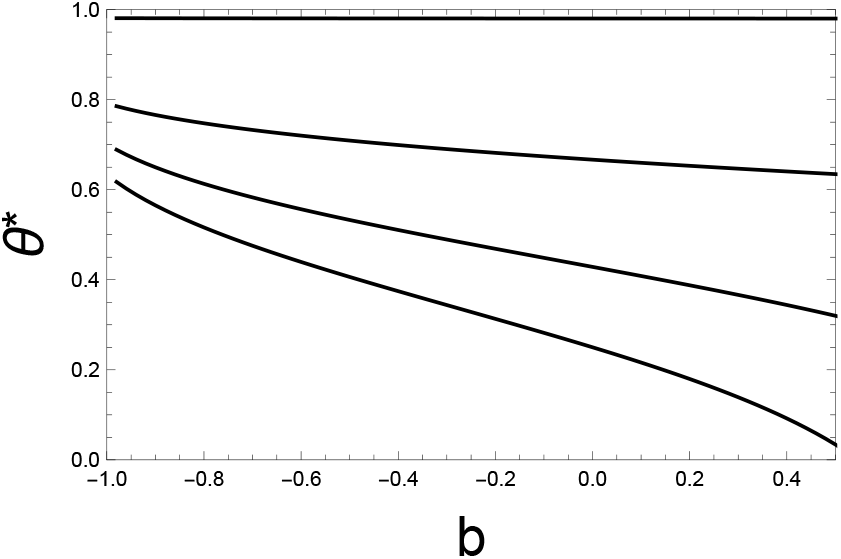
Each curve shows, for the Moran process analyzed in Box 2 with individual fitness (B-i), the singular Kantian coefficient *θ*\* under complete information and incomplete plasticity, for the linear quadratric fecundity function (44), as a function of parameter *b* in that function for *a* = 0.1 and *c* = 1. Each of the four lines corresponds to a different value of the “backward migration probability”, which depends on the exogenously given migration probability *m* (see eq. (B-l) and the description following it). Starting from the top, the first line, where the Kantian coefficient remains essentially constant at *θ*\* = 0.98 is for *m*_b_ = 0.01 whereby *r* = (1 − *m*_b_)*/*(1 + *m*_b_) ≈ 0.98; the second line is for *m*_b_ = 0.2 whereby *r* = (1 − *m*_b_)*/*(1 + *m*_b_) ≈ 0.66; the third line for *m*_b_ = 0.4 whereby *r* = (1 − *m*_b_)*/*(1 + *m*_b_) ≈ 0.42; and the last line, where the Kantian coefficient varies over the range [0, 0.6], is for *m*_b_ = 0.6 whereby *r* = (1 − *m*_b_)*/*(1 + *m*_b_) = 0.25. By computing the Jacobian (A-10) and the Hessian (A-12) coefficients at these values we checked that all these singular Kantian coefficients are indeed both convergence stable and uninvadable.

### 3.3 Complete information and plasticity

#### 3.3.1 Behavioral equilibrium

The defining assumption of our complete information with complete plasticity scenario is that individuals can not only observe the type of the interaction partner but also condition their preferences on it. Hence, the preferences applied in the interaction become state-specific on the interaction. Specifically, we assume that the type *θ* = (*θ*_d_, *θ*_s_) of an individual is a two-dimensional quantitative trait (*θ* ∈ [0, 1]^2^) such that *θ*_s_ parametrizes an individual’s preference (still given by eq. (11)) when individuals in a pair have the same type and *θ*_d_ parametrizes an individual’s preference when individuals in a pair have different types.

In terms of the equilibrium strategies, consider some resident type *θ* = (*θ*_d_, *θ*_s_) and some mutant type *τ* = (*τ*_d_, *τ*_s_) that is different from *θ* (either because *θ*_d_≠*τ*_d_, or because *θ*_s_ ≠ *τ*_s_, or because both *θ*_d_ ≠ *τ*_d_ and *θ*_s_ ≠ *τ*_s_). Then, a resident individual applies the Kantian coefficient *θ*_s_ when interacting with another resident, in which case they both play the equilibrium strategy 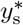, satisfying

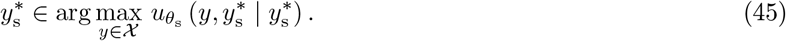

The solution thereof defines the equilibrium strategy as a function of only *θ*_s_ and we write 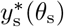 when this dependence needs to be made explicit. In mutant-mutant interactions both individuals apply the Kantian coefficient *τ*_s_ and they both use the equilibrium strategy 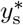, satisfying

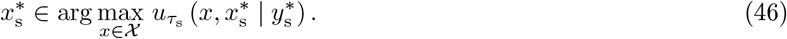

This defines the equilibrium strategy as a function of both *τ*_s_ and *θ*_s_ (since 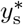 depends on *θ*_s_), which we write 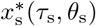. Finally, in mutant-resident pairs where *τ* ≠ *θ*, the mutant applies the Kantian coefficient *τ*_d_ while the resident applies the Kantian coefficient *θ*_d_, and the Nash equilibrium strategies 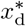 and 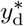 are best responses to each other according to these preferences:

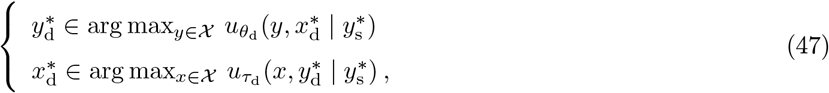

which leads to the dependence of both equilibrium strategies on *τ*_d_, *θ*_d_, and *θ*_s_ and we write 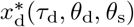 and 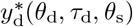. In mutant-resident pairs with the same type, which occurs if individuals from different lineage interact but are of the same type *τ* = *θ*, then each individual in the pair applies the Kantian coefficient *θ*_s_ and therefore expresses strategy 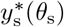.

#### 3.3.2 Evolutionary equilibrium

The fixed point eqs (45)–(47) define the behavioral mechanisms (8)–(10) under complete information with complete plasticity and it follows that the invasion fitness can be written as

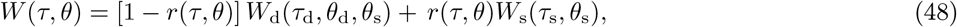

where

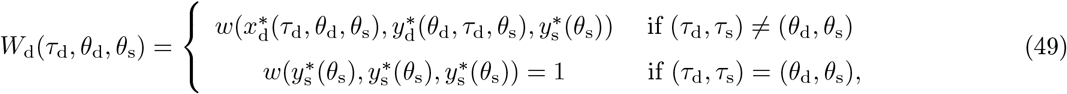

and

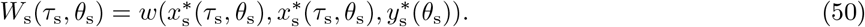

Each component of the mutant trait *τ* = (*τ*_d_, *τ*_s_) thus affects invasion fitness differently. The first trait component, *τ*_d_, only affects the part of invasion fitness *W*_d_ emanating from the interaction of a mutant with a resident. Invasion fitness is not differentiable in trait component *τ*_d_ because, as emphasized by eq. (49), there is generally a discrete jump in the equilibrium strategies at *τ*_d_ = *θ*_d_, from 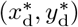 to 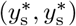) since as long as *τ* ≠ *θ*, individuals in mutant-resident pairs play the equilibrium 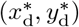, while at *τ* = *θ* the pair plays 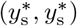 [see the example below for an illustration]. The second trait component, *τ*_s_, not only affects the part of invasion fitness *W*_s_ emanating from the interaction of a mutant with another mutant, but also *W*_d_, since if *τ*_d_ = *θ*_d_ the resident applies the value *θ*_d_ when interacting with a mutant if *τ*_s_ ≠ *θ*_s_ but instead the value *θ*_s_ if *τ*_s_ = *θ*_s_. The invasion fitness (48) is thus not differentiable in the mutant type (*τ*_d_, *τ*_s_). Moreover, as will be shown and explained in detail in an example below, there is typically an infinite number of uninvadable types *θ* ∈ [0, 1]^2^. In spite of these challenges, we have identified two simple sufficient conditions for a type to be uninvadable (but we were unable to conclude on convergence stability).

To state our result, for any *θ*_d_, *τ*_d_, *θ*_s_ ∈ [0, 1] define

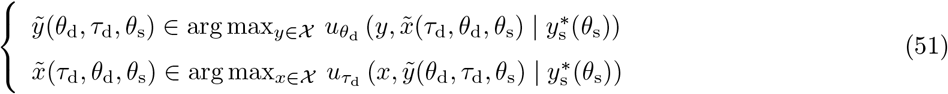

and

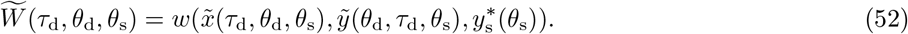

##### Result 3.

*A sufficient condition for W*_d_(*τ*_d_, *θ*_d_, *θ*_s_) ≤ 1 *for all τ*_d_ ∈ [0, 1] *and W*_s_(*τ*_s_, *θ*_s_) ≤ 1 *for all τ*_s_ ∈ [0, 1] *(and thus W* (*τ, θ*) ≤ 1 *for all τ* = (*τ*_d_, *τ*_s_) ∈ [0, 1]^2^*) is that θ* = (*θ*_d_, *θ*_s_) = 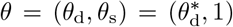, *where* 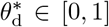 *is such that* 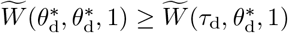 *for all τ*_d_ ∈ [0, 1].

For a formal proof of this result, see Appendix C. To understand this result, note first that *θ*_s_ = 1 guarantees that a mutant cannot achieve a higher fitness when interacting with another mutant, than a resident does when interacting with another resident. This value of the Kantian coefficient indeed implies that both individuals in the interaction act as “social planners”: they choose that strategy which, when chosen by both individuals, maximizes their fitness. Second, note that for *τ*_d_ ≠ *θ*_d_, the system of equations (51) is mathematically equivalent to the system of equations that characterizes the equilibrium strategies between a resident and a mutant (see (47)), and 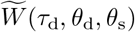 is equivalent to *W*_d_(*τ*_d_, *θ*_d_, *θ*_s_).^11^ Hence, the value 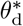 defined in the result is such that if residents apply 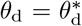 when interacting with mutants, the residents get a higher fitness than the mutants do in mutant-resident interactions, for any value of *τ*_d_ ≠ *θ*_d_. Thus, 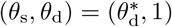 is a “safe bet” as an uninvadable point.

However, and as mentioned before, there are typically many other uninvadable types. To see why, it is useful to draw a parallel between the function 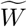, defined in eq. (52), and the fitness of a mutant under the complete information and incomplete plasticity scenario when relatedness is equal to zero, defined in (38). These are mathematically equivalent, except that in eq. (52) individuals in the resident population play strategy 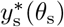, where *θ*_s_ may differ from *θ*_d_. This parallel allows us to realize that the same arguments as for the derivation of Result 2 can be applied here, to conclude that a relevant candidate value for 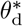 is the one satisfying 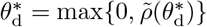 where

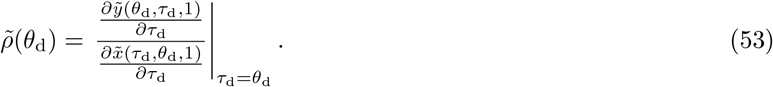

From (differentiable) fitness 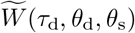 and following the same arguments as those leading up to Result 2, we know that an uninvadable type satisfying 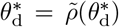 cannot be equal to one, i.e., 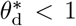. Based on the same argument as used in the previous paragraph, we have 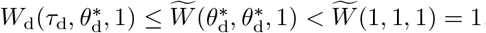. The inequality 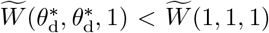 follows from the fact that *θ*_s_ = 1 implies that both individuals in the interaction act as “social planners” and thus maximize their fitness, and hence interaction partners with 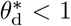 do *not* act as social planners, thereby choosing a strategy that does not maximise their fitness (see also Appendix C).

We can thus conclude that 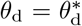 implies that *W*_d_(*τ, θ*) <1 for any 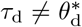. Hence, we can further conclude that 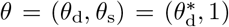 implies that *W* (*τ, θ*) <1 for any *τ* ≠*θ*. It is this discrete jump between the invasion fitness of any mutant *τ* ≠ *θ* and the residents’ fitness that opens the door for the multiplicity of uninvadable types. For example 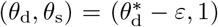 for some sufficiently small *ε* > 0 would also be uninvadable: with this resident type, a mutant can achieve a higher fitness than a resident in a mutant-resident interaction, but one can always find *ε* small enough so that this advantage would be too small to overcome the difference 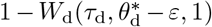. Likewise, 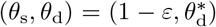 for some sufficiently small *ε >* 0 would also be uninvadable: with this resident type, mutants can achieve a higher fitness when interacting with each other than when two residents interact, but one can always find *ε* small enough so that this advantage would be too small to overcome the difference 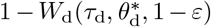. Note that this example further shows that *θ*_s_ = 1 is not necessary for *θ* to be uninvadable.

In order to illustrate Result 3, we work out an example assuming a constant relatedness of 1*/*2 and an individual fitness function of the form *w*(*x*_*i*_, *x*_*−i*_, *y*) = *f* (*x*_*i*_, *x*_*−i*_)*/f* (*y, y*), with fecundity 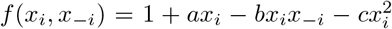 given by the linear-quadratic function (44). This model could be thought of as interactions between siblings in a family-structured semelparous population. Given these assumptions, the invasion fitness (48) is

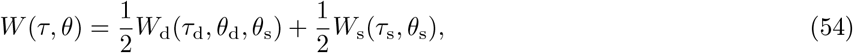

with

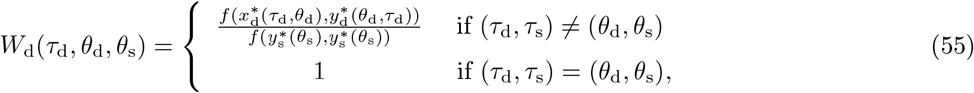

and

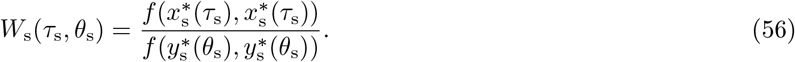

In force of eq. (45), the (Nash) equilibrium strategy for a resident pair is

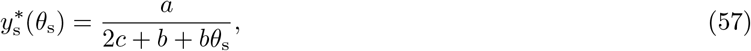

while in force of eq. (47), the strategies for a mutant-resident pair are\

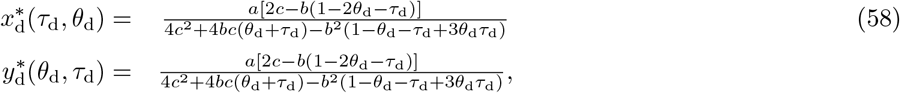

and in force of eq. (46), the equilibrium strategy for a mutant pair is

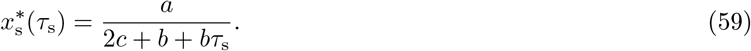

For this model, we can also write for any *τ*_d_, *θ*_d_ ∈ [0, 1] that

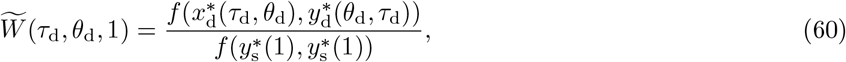

and the candidate uninvadable Kantian coefficient thus satisfies

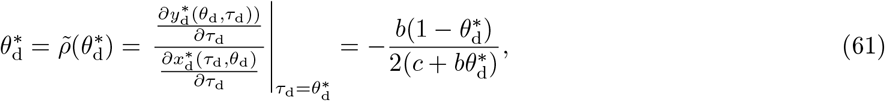

which has a relevant root in the interval [0, 1] when *b* <0 (which implies that the strategies are strategic complements), given by

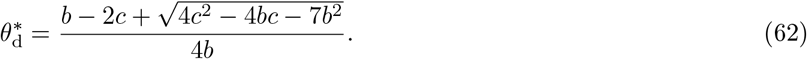

It is straightforward to check, for instance by computing 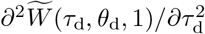 at 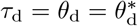, that there is a range of parameter values where this Kantian coefficient is uninvadable and this is illustrated in panel A of Fig. 2.

**Figure 2:**
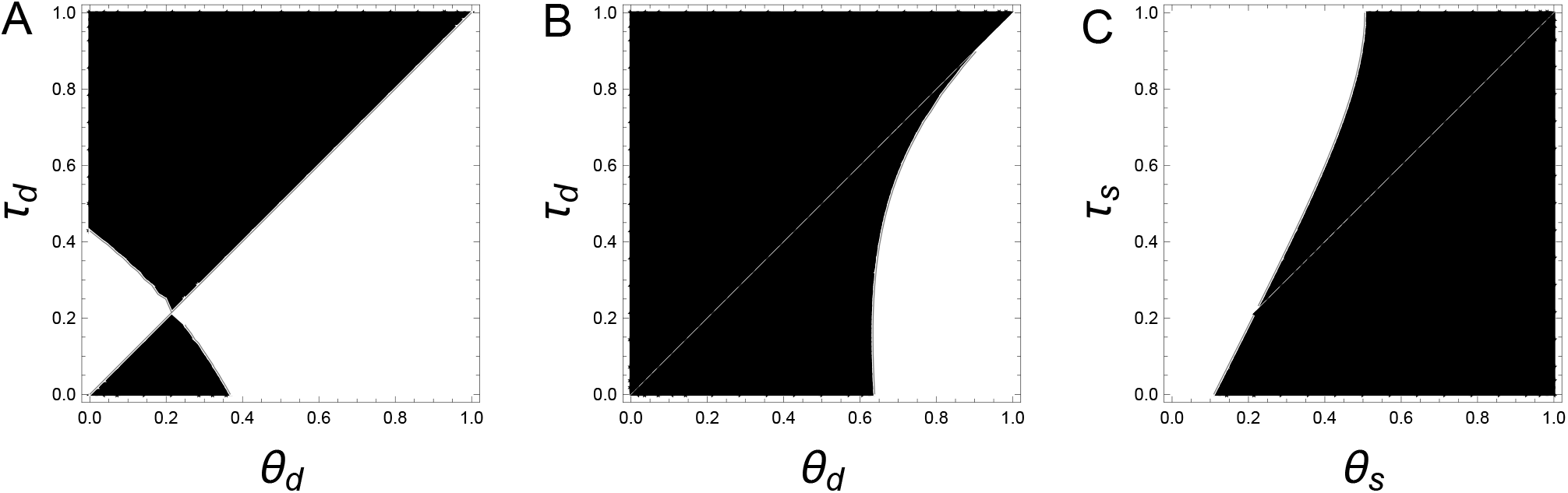
Pairwise invasibility plots. **Panel *A*** *displays the difference* 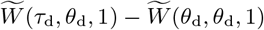 for all combinations of *θ*_d_ (*x*-axis) and *τ*_d_ (*y*-axis) values (and thus holding *θ*_s_ = 1 fixed), for the parameter values *a* = 0.1, *b* = −0.5 and *c* = 1 of the linear quadratic game (44) 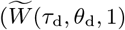 is thus defined by (60). The difference is zero on the diagonal, since *τ*_d_ = *θ*_d_. The black region depicts all the combinations (*θ*_d_, *τ*_d_) such that the difference is negative, while the white region outside the diagonal depicts all the combinations (*θ*_d_, *τ*_d_) such that the difference is strictly positive. The candidate given by eq. (62) is 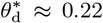, which the graph shows is uninvadable. **Panel B** displays the difference *W* (*τ, θ*) − *W* (*θ, θ*) determined by eqs. (54)–(59) of a mutant *τ* = (*τ*_d_, 1) in a resident *θ* = (*θ*_d_, 1) population for all combinations of resident *θ*_d_ (*x*-axis) and mutant *τ*_d_ (*y*-axis) trait values (and thus holding *θ*_s_ = 1 and *τ*_s_ = 1 fixed) for the parameter values *a* = 0.1 *b* = −0.5 and *c* = 1 of the linear quadratic game (44). On the diagonal, the difference equals zero (and invasion fitness equals one) since *τ*_d_ = *θ*_d_. The black region depicts all combinations (*θ*_d_, *τ*_d_) such that the difference is negative, i.e., *W* (*τ, θ*) <1, so that the mutant *τ*_d_ cannot invade, while the white region outside the diagonal depicts all combinations (*θ*_d_, *τ*_d_) such that the difference is positive, *W* (*τ, θ*) *>* 1, so that the mutant can invade. The graph thus shows that all values of (*θ*_d_, 1) with *θ*_d_ below approximately 0.6 are uninvadable. There is thus a multiplicity of uninvadable types, which necessarily contains that of Panel A. Note that resident values above approximately *θ*_d_ ≈ 0.9 are invadable by any mutation *τ*_d_ <*θ*_d_, while resident values between approximatively 0.6 and 0.9 are invadable by mutants with *τ*_d_ = *θ*_d_ − δ for δ *>* 0 large enough. **Panel C** displays *W* (*τ, θ*) − *W* (*θ, θ*) determined by eqs. (54)–(59) for a mutant *τ* = (*τ*_d_, *τ*_s_) in a resident *θ* = (*θ*_d_, *θ*_s_) population for all combinations of resident *θ*_s_ (*x*-axis) and mutant *τ*_s_ (*y*-axis) trait values holding 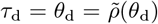 fixed and given by eq. (61) under the parameter values *a* = 0.1 *b* = −0.5 and *c* = 1 of the linear quadratic game (44). As for Panel B, the black region depicts all combinations (*θ* ;_s_, *τ* ;_s_) such that the difference is negative, i.e., *W* (*τ, θ*) <1, so that the mutant *τ*s cannot invade, while the white region outside the diagonal depicts all combinations (*θ* ;_s_, *τ* ;_s_) such that the difference is positive, *W* (*τ, θ*) *>* 1, so that the mutant can invade. The graph thus shows that all values of 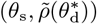 with *θ*_s_ above approximately 0.5 are uninvadable.

This example illustrates the two features of Result 3 discussed in general terms above. First, as long as *τ* ≠ *θ*, individuals in mutant-resident pairs play the equilibrium (58), while at *τ* = *θ*, they play eq. (57). This makes the strategies discontinuous at *τ*_d_ = *θ*_d_, which in turn implies that invasion fitness is not differentiable. Despite this discontinuity embedded in fitness *W*_d_(*τ*_d_, *θ*_d_, *θ*_s_) (eq. (55)), Result 3 shows that we can use the function 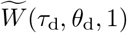 to straightforwardly calculate 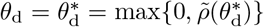, and then use the Hessian to check that 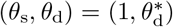 is indeed uninvadable. This is illustrated in panel A of Fig. 2. Second, there are multiple uninvadable types; this is illustrated in panels B and C of Fig. 2.

Our formalization of preference evolution under complete information and plasticity makes it difficult to reach any specific conclusion about long term evolution for two reasons. First, because invasion fitness is not differentiable, a different toolkit than the usual multidimensional convergence stability criterion is needed to characterize the attractor points of the evolutionary dynamic (Leimar, 2009). Second, the multiplicity of uninvadable equilibria compounded with the non differentiability makes the focus on monomorphic population questionable and a treatment with polymorphic populations appears required. It would thus be relevant to analyze a full dynamics model of preference evolution with mutation and selection under complete information and plasticity to reach more definite results and also to determine whether the equilibrium 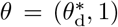 identified in Result 3 plays a special role, for instance whether it is a stochastically stable trait value (*sensu* Foster and Young, 1990).

## 4 Discussion

By investigating the evolution of semi-Kantian preferences under different informational and behavioral plasticity assumptions in group-structured populations, we have extended the evolutionary viability analysis of this class of preferences. While we restricted attention to pairwise interactions, and preferences characterized by a single evolving quantitative trait, the Kantian coefficient, our model weaves together different threads of the literature and shows how long-term evolution concepts can be used to analyze preferences under gradual evolution. We obtained three main results on the convergence stability and uninvadability of the value of the Kantian coefficient.

First, when interacting individuals have no information about each other’s Kantian coefficient and mutants hold beliefs about the probability of being matched with another randomly sampled mutant from the same lineage, we confirm that an uninvadable Kantian coefficient must equal the coefficient of relatedness (Alger and Weibull, 2013; Alger et al., 2020). But instead of considering the set of possible utility functions to be the set of all continuous functions as done in this previous work, we focused on the more restricted setting where utility functions are parametrized by a single quantitative trait. This allowed us to cover not only uninvadability in a complementary and less abstract way, but also to cover convergence stability. In Result 1, we show that the Kantian coefficient equal to the coefficient of relatedness is both convergence stable and uninvadable when trait effects on relatedness are sufficiently weak. Thus, we characterize conditions where gradual evolution drives preferences to induce individuals to behave according to Hamilton’s (marginal) rule at the strategy level. One relevant avenue for future research for preference evolution under incomplete information is to consider more realistic demographic scenarios of class structured population (e.g., by sex, age, or stage).

Second, when interacting individuals can observe each other’s type, but each individual has the same preferences regardless of the other’s type, we showed that an uninvadable value of the Kantian coefficient can exceed, fall short of, or equal the coefficient of relatedness. Moreover, we showed that the sign of the discrepancy is determined by whether an individual’s equilibrium strategy is correlated positively, negatively, or not at all with the opponent’s equilibrium strategy. This response coefficient in turn depends on the specifics of the individual fitness function so that the Kantian coefficient will depend on life-history parameters such as survival, migration, etc. [recall the example of the fitness function (B-i)]. Gradual evolution thus now drives preferences to induce individuals to behave according to a context-specific Kantian coefficient, which combines both the relatedness and the response coefficients. This result is fully in line with previous models under complete information and incomplete plasticity, which have all considered other parametric classes of preferences than the one we examined (e.g. Bester and Güth, 1998; Bolle, 2000; Possajennikov, 2000; Heifetz et al., 2007b,a; Akçay and Van Cleve, 2009 for models without relatedness, and Alger, 2010; Alger and Weibull, 2010, 2012; Akçay and Van Cleve, 2012 for models with relatedness). The dependence of an uninvadable value of the Kantian coefficient on the response coefficient stems from the commitment to a particular behavioral response that an individual’s preferences induces. By being observable, a mutant’s preference type can thus induce a resident to adopt a different strategy than the one adopted in an interaction with another resident, an effect that is absent under incomplete information. Although we established necessary conditions for a Kantian coefficient value to be uninvadable and illustrated uninvadablility and convergence stability under a linear quadratic game (Fig. (1)), we did not succeed in identifying simple general sufficient conditions, neither for uninvadability nor for convergence stability. In particular, we cannot rule out *evolutionary branching points*, which obtain when a singular Kantian coefficient value is convergence stable but not uninvadable (recall footnote 3, and see Geritz et al., 1998 for a general discussion and McNamara and Leimar, 2020 for typical evolutionary game theory applications). An avenue for future research on preference evolution under complete information is thus to analyze conditions leading to polymorphism in preferences. ^12^

Finally, we considered the case of complete information with complete plasticity where individuals can both observe the opponent’s type and also condition its preferences on it. This is akin to a green-beard or secret handshake mechanism (Hamilton, 1964b; Grafen, 1990; Robson, 1990), but at the preference level rather than at the strategy level as in most previous work. Since an individual’s Kantian coefficient may depend on the type of the interaction partner, a type is now a two-dimensional quantitative trait. Compared to the complete information incomplete plasticity scenario, individuals are thus no longer committed to respond according to one and the same Kantian coefficient. Residents are therefore less exploitable by mutants and individuals can be regarded as implementing multiple selves (Lester, 2015) since their preferences and motivations are context-dependent. As we showed, this implies that residents can be pure Kantians when interacting with each other, and still be uninvadable: they can prevent entry by mutants by using the Kantian coefficient equal to the response coefficient when interacting with individuals with a different type than theirs. In such a population, when interacting with each other residents then use the strategy which yields the highest possible individual fitness. In other words, they use the strategy that yields an efficient outcome. This is reminiscent of a result by Dekel et al. (2007), who showed that a class of “coordination” preferences, which results in efficient strategy profiles, are stable. It is also reminiscent of results obtained by Wang and Wu (2023) in a model where the search and matching of interacting partners is endogenous but costless: the authors demonstrate the evolutionary stability of a preference for interacting with an individual with the same type as self combined with play of an efficient strategy profiles. We were, however, were not able to characterize convergence stability nor the Kantian coefficient when individuals interact with others with a type different from their own type. Ascertaining this in the non-differentiable setting of complete information with complete plasticity is thus left for future work.

We derived these three results assuming existence of a unique behavioral equilibrium in the resident population and when mutant-resident (or mutant-mutant) interactions occur. Given that it is common that interactions produce a multiplicity of (Nash) equilibria, an important avenue for future research will be to examine the robustness of our results in settings with multiple equilibria. Several questions will need to be addressed. First, one can imagine several alternative formalizations of the interaction between individuals when multiple equilibria are possible. For instance, one could assume that different pairs of individuals in the resident population play different equilibria; such a model would likely require tools from the literature on class-structured populations (since an average individual may then play different equilibria and thus be in different contexts or states, the defining feature of class structured populations e.g., Avila and Mullon, 2023). An alternative would be to assume that the same equilibrium is played in all resident-resident interactions; our formalization should then apply to each equilibrium separately (with implications for possibly different relatedness coefficients depending on the equilibrium being played). The same modeling choices will arise when there exist multiple equilibria in mutant-resident pairs. Second, the definitions of uninvadability and convergence stability may need to be adapted. In particular, for uninvadability, would a preference type be deemed uninvadable only if it is uninvadable for all possible formalizations of interactions in the presence of multiple equilibria? A similar question arises for convergence stability. For existing attempts to address some of these questions, see Ok and Vega-Redondo (2001); Dekel et al. (2007); Alger and Weibull (2013, 2016); Alger et al. (2020); Alger and Weibull (2023); Wang and Wu (2023).

Our three results show within the same model how different information and behavioral flexibility assumptions lead to different values of the Kantian coefficient, and thus to different equilibrium strategies. By contrast to strategy evolution models, which predict behavioral patterns for stationary environments, preference evolution models allow to make predictions about behavioral change in a new environment, and can be tested, using either field data or experimental data. A key prediction of our model is that equilibrium behavior at evolutionary equilibrium should be in accordance with Hamilton’s marginal rule expressed at the strategy level only if evolution operated on interactions under incomplete information (recall equation (26)), and that if observed deviations from this rule resulted from interactions taking place under complete information, the deviation from Hamilton’s marginal rule should depend on the specifics of the interaction at hand (recall equation (43)). In some experiments under incomplete information, humans do appear to conform to behave according to Hamilton’s (marginal) rule at the strategy level (Levy and Lo, 2022).

Economic games can further be used to discriminate between different preferences in controlled laboratory experiments. For example, Fisman et al. (2007) and Bruhin et al. (2019) use dictator games to estimate individuals’ preference parameters assuming other-regarding preferences. More recent studies in this literature have been inspired by results in the evolutionary theory of preferences to design economic games that further allow the researchers to discriminate between other-regarding and semi-Kantian preferences (Miettinen et al., 2020), and to estimate the importance of Kantian concerns relative to other-regarding concerns (Van Leeuwen and Alger, 2022). The latter study indicates that many individuals appear to be driven by a combination of self-interest, semi-Kantian concerns, and other-regard. This finding is in line with evolutionary arguments showing that it is important to distinguish between preferences expressed in terms of fitness consequences and preferences expressed in terms of payoff consequences (Lehmann et al., 2015; Alger et al., 2020). The qualitative nature of evolved preferences at the material payoff level indeed differ from that at the fitness level, and this difference depends on demographic properties under genetic evolution and transmission rules under cultural evolution (Alger et al., 2020). Payoff and fitness incentives tend to agree in panmictic and family-structured populations, but tend to disagree in spatially structured populations owing to the presence of local competition between individuals. Future research could adopt a similar approach to the one we develop here, by analyzing evolution of preferences expressed at the level of material payoffs under a variety of informational assumptions.

In this paper we have sought to bring together modeling tools from evolutionary biology and from economics in order to investigate the evolution of preferences that guide behavior in strategic interactions. We hope that our formalization has illustrated some of the nuances, intricacies, and richness of research endeavours that seek to analyse more completely the evolutionary dynamics of behavioral mechanisms, and that it will inspire future studies into the many open theoretical and empirical research questions.

## Declarations

## Author’s contributions

L.L. conceived the model, I.A. and L.L. derived the results, L.L. wrote the first draft and both authors finalized the manuscript.

## Funding

I.A. acknowledges IAST funding from the French National Research Agency (ANR) under grant ANR-17-EURE-0010 (Investissements d’Avenir program), and funding from the European Research Council (ERC) under the European Union’s Horizon 2020 research and innovation programme (grant agreement No 789111 - ERC Evolving Economics).

### Box 1.

Invasion fitness as eigenvalue

The invasion fitness of a type is its geometric growth ratio when rare in a resident population (Fisher, 1930; Eshel and Feldman, 1984; Metz et al., 1992; Ferrière and Gatto, 1995). When the resident population is monomorphic for *θ*, the invasion fitness *W* (*τ, θ*) of mutant *τ* under our demographic assumptions (section (2.1)) is obtained as the leading eigenvalue of the matrix

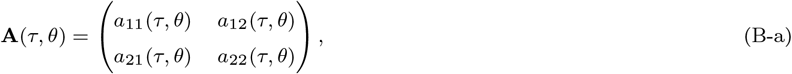

where *a*_*ij*_ stands for the expected number of groups with *i* ∈ {1, 2} mutants that over one demographic time period descend (either through local change or through migration) from a focal group with *i* ∈ {1, 2} mutants, when the population is otherwise monomorphic for *θ*. Matrix **A**(*τ, θ*) is assumed to be regular (irreducible and aperiodic, Iosifescu, 2007, p. 123). It then follows from standard results on multitype branching processes that the lineage of a single *τ* mutant goes extinct with probability one if, and only if, *W* (*τ, θ*) ≤ 1, otherwise the lineage spreads into the population when rare and becomes infinitely large (Harris, 1963; Karlin and Taylor, 1975). By definition of invasion fitness, *W* (*τ, θ*)**u**(*τ, θ*) = **A**(*τ, θ*)**u**(*τ, θ*), where **u**(*τ, θ*) = (*u*_1_(*τ, θ*), *u*_2_(*τ, θ*)) is the only non-negative right eigenvector of **A**(*τ, θ*), where, by normalization, *u*_1_(*τ, θ*) + *u*_2_(*τ, θ*) = 1. The eigenvector **u**(*τ, θ*) can be interpreted as the quasistationary distribution of mutant group types as it is invariant to multiplication by **A**(*τ, θ*), whereby *u*_*i*_(*τ, θ*) is the frequency of groups with *i* ∈ {1, 2} mutants among groups with at least one mutant. Following previous developments (Mullon et al., 2016), we can left multiply *W* (*τ, θ*)**u**(*τ, θ*) = **A**(*τ, θ*)**u**(*τ, θ*) by the vector (1, 2). Rearranging terms, this produces

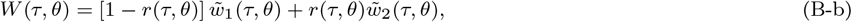

where

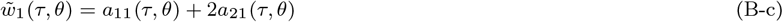

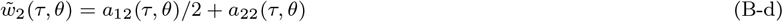

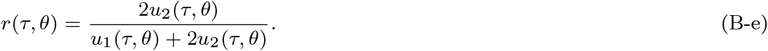

Here, 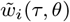 is the expected total number of individuals produced (including the surviving self) by a single *τ* individual over one demographic time step when there are *j* ∈ {1, 2} *τ* individuals in its group and the population is otherwise monomorphic for *θ*; and *r*(*τ, θ*) is the probability that, for any given descendant of the initial mutant, the neighbor of that mutant is also a mutant. An explicit example of these invasion fitness components is given in Box 2.

Eq (B-b) is a recipient-centered representation of the mutant’s geometric growth ratio since it is expressed as the average of the expected fitness of a type *τ* individual, who is necessarily the recipient of the traits of others. An actor-centered representation of the growth ratio, which focuses on the consequence on others of an individual expressing the mutant instead of the resident trait value can also be obtained (Hamilton, 1970; Rousset, 2015). Such an actor-centered representation of invasion fitness can be reached by rearranging the components of eq. (B-b) (Lehmann and Rousset, 2020). Indeed, owing to the fact that 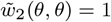, we have the equality

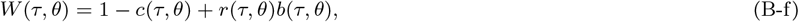

where

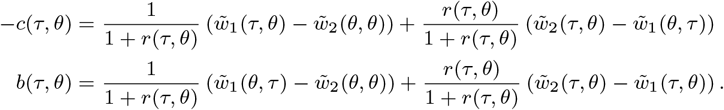

Here, *−c*(*τ, θ*) is the *average effect* (sensu Fisher, 1941) on the number of mutant gene copies produced by a single individual when expressing a copy of the mutant instead of the resident allele. The average thus being over the two possible contexts in which an individual expressing *τ* instead of *θ* can be: interacting with a neighbor that carries or not the mutant. The actor-centered perspective of eq. (B-f) is then born out from the fact that *b*(*τ, θ*) is the average effect on the expected number of offspring produced by an individual’s neighbour, which stemming from the actor switching to expressing a copy of the mutant instead of the resident allele.

### Box 2.

Moran process example

We illustrate the invasion fitness components described in Box 1 by considering a process where exactly one individual dies in each group during a demographic time step (i.e., an instance of a Moran process, Moran, 1962). For this case, the entries of matrix (B-a) are

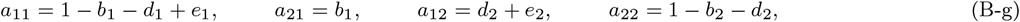

with *b*_*i*_ and *d*_*i*_ standing, respectively, for the probability that there is a mutant descendant and mutant death, and *e*_*i*_ is the expected number of succesful emigrant mutants, in a group with *i* mutants. These variables are given by

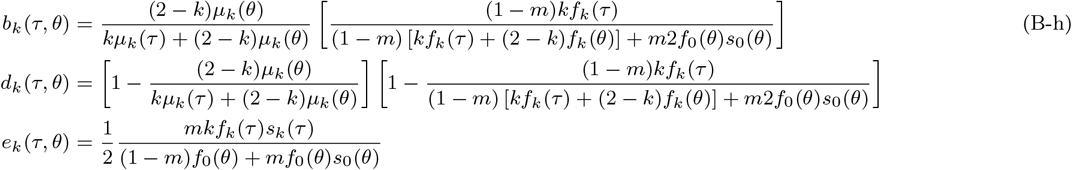

where *f*_*k*_(*θ*^*′*^), *μ*_*k*_(*θ*^*′*^), *s*_*k*_(*θ*^*′*^) are, respectively, the fecundity, death-factor, juveniles’ survival probability during migration, of a single type *θ*^*′*^ ∈ {*τ, θ*} adult individual when there are exactly *k* mutants in its group (see Lehmann et al., 2015; Mullon et al., 2016 for more details on the derivation and the case where there are more than 2 individuals per group). On setting 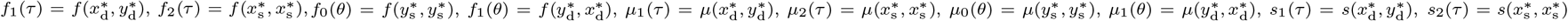, and 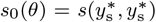, where *x* refers to mutant and *y* to resident strategies [recall eqs. (6)–(7)] and *f* : *χ* ^2^ → ℝ_+_, *μ* : *χ* ^2^ → ℝ _+_, and *μ* : *χ* ^2^ → ℝ _+_, then algebraic rearrangements show that the fitness function *w* : *χ* ^3^ → ℝ_+_ in eqs. (6)–(7) for the Moran process is defined as

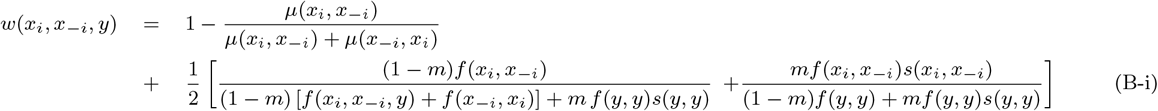

(see Box 1 of Lehmann et al., 2015 for a biological interpretation of each term).

Even for this Moran process, the expression for relatedness eq. (B-e) is complicated, but its computation can be alleviated by using an invasion fitness proxy. An invasion fitness proxy is by definition any fitness measure *P* (*τ, θ*) that is sign equivalent to *W* (*τ, θ*) such that the evolutionary invasion analysis can be carried out from this measure (i.e. *P* (*τ, θ*) ≤ 1 ⟺ *W* (*τ, θ*) ≤ 1). An invasion fitness proxy for *W* (*τ, θ*) can be obtained by keeping the functional form eq. (B-b), but relatedness, instead of being given by the complicated expression eq. (B-e), is given by

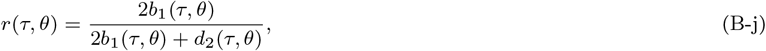

which can be readily evaluated using eq. (B-h). Conceptually, this simplification obtains by substituting *u*_*i*_ → *t*_*i*_ in eq. (B-e), where *t*_*i*_ is the sojourn time with *i* ∈ {1, 2} mutants of the mutant lineage in a single group where *t*_1_ = 1*/d*_1_ and *t*_2_ = *b*_1_*/*(*d*_1_*d*_2_) (see Lehmann et al., 2015; Mullon et al., 2016 for more details). Substituting eq. (B-h) into eq. (B-j) and using the expression for the vital rates in terms of strategies and assuming, for simplicity that fecundity *f* is independent of the types, one can then check that relatedness can be written as

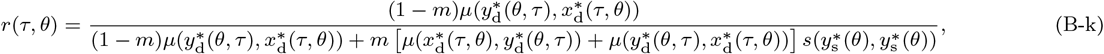

where we made explicit all functional dependencies. Further, in a monomorphic population relatedness boils down to

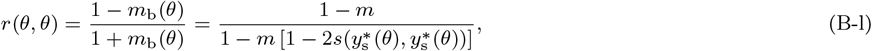

where 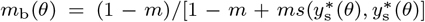 is the backward migration probability, i.e., the probability that an individual randomly sampled in a patch is of philopatric origin. Eq. (B-l) displays two generic features about relatedness. First, it is a monotonic decreasing function of dispersal and of juvenile survival. Second, relatedness can depend endogeneously on the interactions, because the spatial structure is an outcome of survival and reproduction, which are themselves functions of interactions between individuals. If survival 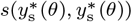 were independent of strategies, then neutral relatedness would be independent of the types and reduce to *r* = (1 − *m*)*/*(1 − *m*(1 − 2*s*), as it should (Mullon et al., 2016), for parameters *m* ∈ (0, 1] and *s* ∈ [0, 1].

## Appendix A Behavioral perturbation under incomplete information

We here show that the behavioral perturbation (18) is always positive, i.e., *∂x*\* (*τ, θ*)*/∂τ* |_*τ*=*θ*_ *>* 0. We proceed in two steps. First, we evaluate the perturbation of a mutant’s reaction function and show that this is positive. Second, we show that this latter perturbation is sign equivalent to *∂x*\* (*τ, θ*)*/∂τ* |_*τ*=*θ*_.

For the first step recall from eq. (13) that a specific mutant’s expected utility is

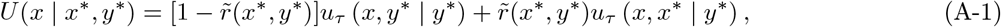

where the notation | *x*\*, *y*\* emphasises that we are here holding the strategies of the other mutant and resident individuals as given. A mutant’s best response to (*x*\*, *y*\*) is implicitly defined by the necessary first-order condition:

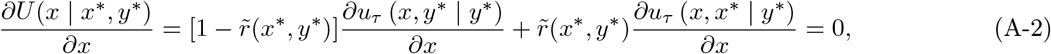

which, using eq. (11), is written

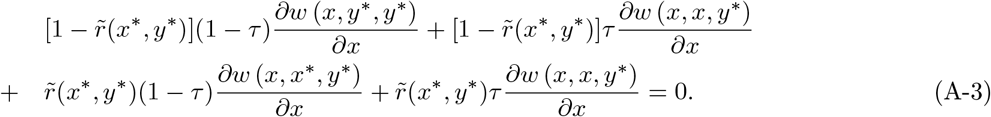

This equation implicitly defines the specific mutant’s optimal strategy *x*(*τ, x*\*, *y*\*) as a function of *τ, x*\*, and *y*\* (which depends on *θ*). In other words, it defines this mutant’s *reaction function*, which specifies its utility-maximizing strategy for each (*x*\*, *y*\*) (e.g., Fudenberg and Tirole, 1991, p. 14). We use an index *i* to denote the partial derivative with respect to the *i*-th argument of the individual fitness function to rewrite eq. (A-3) as follows:

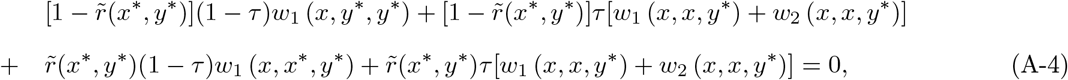

which simplifies to

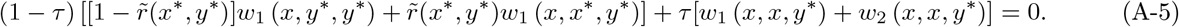

Applying the implicit function theorem holding (*x*\*, *y*\*) fixed, we obtain:

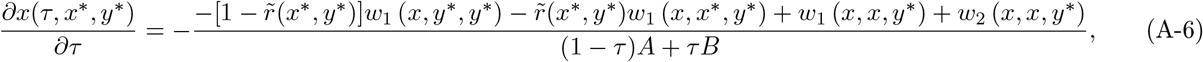

where

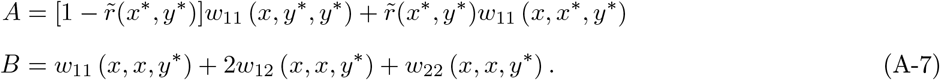

Strict concavity of *u*_*τ*_ for all *τ* ∈ [0, 1] implies that *A* <0 and *B* <0 and so the denominator of eq. (A-6) is strictly negative. At equilibrium, and locally at *τ* = *θ*, we have *x* = *x*\* = *y*\*, implying that the numerator reduces to *w*_2_(*x, x, y*\*), which is strictly positive. Hence, the perturbation *∂x*(*τ, x*\*, *y*\*)*/∂τ* |_*τ*=*θ*_, which measures this mutant’s (infinitesimal) change in its reaction function *x*(*τ, x*\*, *y*\* (*θ*)) when the value of its Kantian coefficient is infinitesimally increased at *τ* = *θ*, is strictly positive.

We turn now to the second step to show why the result derived in the first step implies that the mutant equilibrium strategy *x*\* must increase as a result of an (infinitesimal) increase in the value of *τ* for all mutants. First, note that such an increase in *τ* has no effect on the resident strategy (see eq. (13)), implying that we can indeed hold *y*\* fixed, as we did above. Hence, for a given *τ* and *y*\*, the equilibrium strategy *x*\* (*τ, θ*) is a fixed point of the mutant’s best-response function (see the second line of eq. (13)) and this is a point on the curve described by the reaction function *x*(*τ, x*\*, *y*\*) for any given (*τ, y*\*).. Because *∂x*(*τ, x*\*, *y*\*)*/∂τ* |_*τ*=*θ*_ describes how the response function of one mutant is slightly displaced by the introduction of a small mutant deviation at *τ* = *θ*, and because we proved above that any such deviation is positive for any strategy *x*\* played by other mutants, the equilibrium strategy *x*\* (*τ, θ*) must locally vary with the same sign as the reaction function varies, i.e, *∂x*\* (*τ, θ*)*/∂τ* |_*τ*=*θ*_ ∝ *∂x*(*τ, x*\*, *y*\*)*/∂τ* |_*τ*=*θ*_.

## Appendix B Jacobian and Hessian under complete information with incomplete plasticity

Differentiating eq. (40) with respect to *θ* and evaluating at *θ*\* satisfying *S*(*θ*\*) = 0 yields

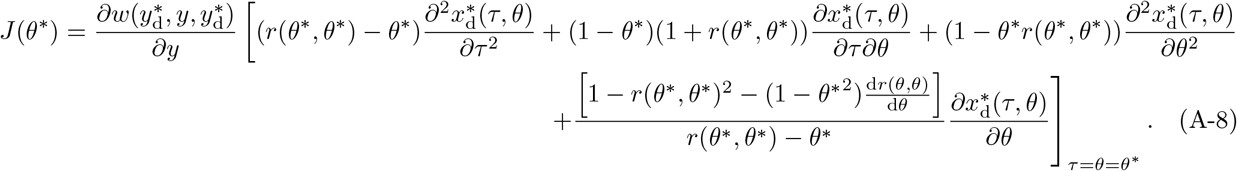

Using eq. (40) at *S*(*θ*\*) = 0 and using eq. (36) we can express the singular trait value implicitly as

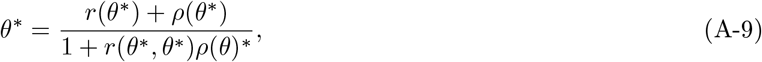

which, on substituting into eq. (A-8), using 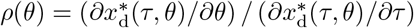 and simplifying produces

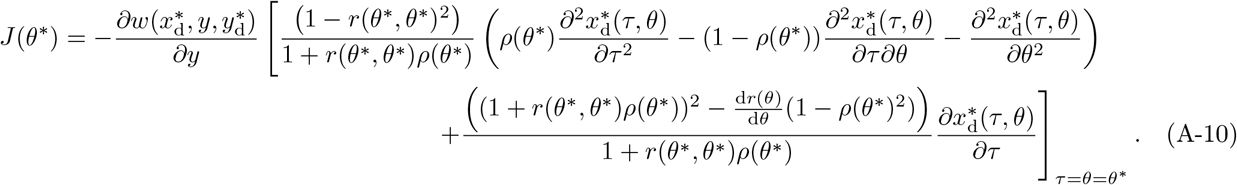

The second order behavioral perturbations 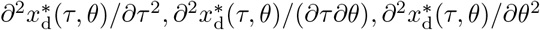 appearing in this Jacobian can be computed by using implicit differentiation in eq. (31). The resulting expressions are complicated and lengthy and we were unable to infer some general information from these expressions, although they can be handled easily with a symbolic manipulation system such as Mathematica (Wolfram Research, 2016). A Mathematica notebook with all algebraic computations of the paper is available on request.

Now using invasion fitness (38) to evaluate *H*(*θ*) = *∂*^2^*W* (*τ, θ*)*/∂τ* ^2^|_*τ*=*θ*_, we find that

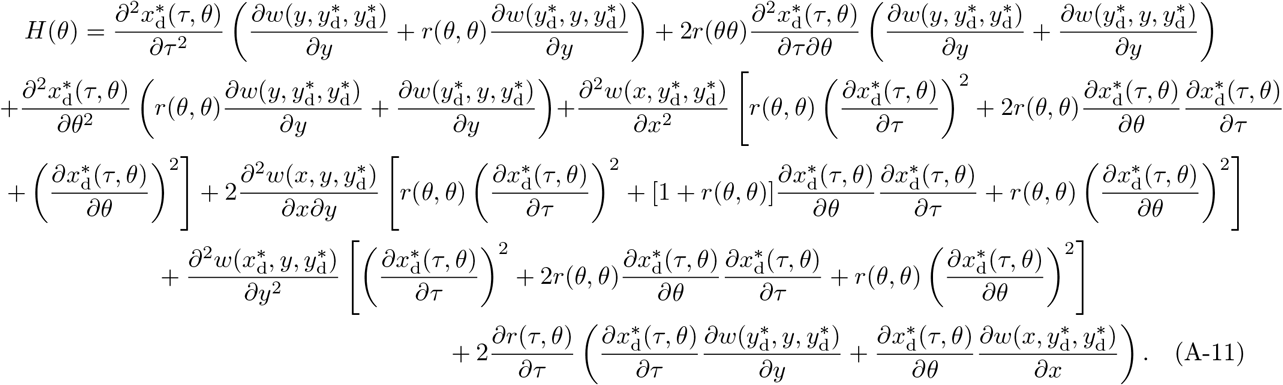

Substituting into this expression 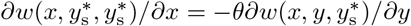 and eq. (A-9) yields

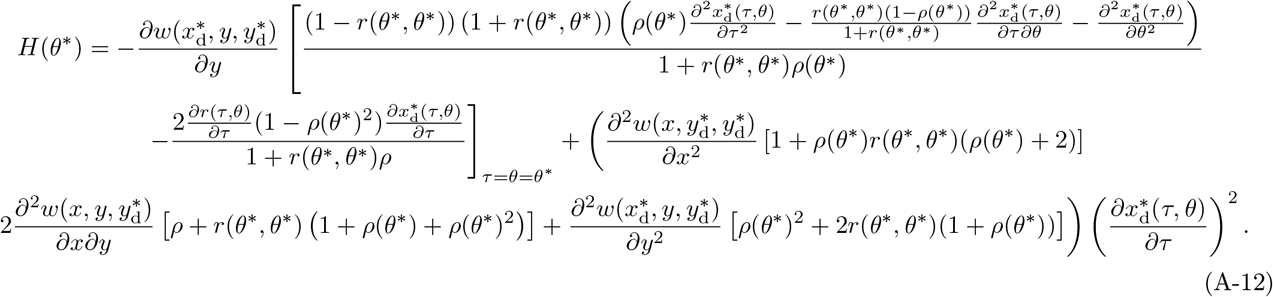

A key distinction between the Jacobian *J* (*θ*\*) and the Hessian *H*(*θ*\*) is that the sign of the Jacobian does not depend directly on fitness derivatives, while the Hessian does. Both expressions remain complicated and we did not manage to obtain general information from them. Hence, they need to be evaluated on a case by case basis.

## Appendix C Proof of Result 3

We first examine mutant-mutant interactions (the term in eq. (50)) and then turn to mutant-resident interactions (the term in eq. (49)).

Consider some resident type *θ* = (*θ*_d_, *θ*_s_) ∈ [0, 1]^2^. For any such type, residents obtain individual fitness 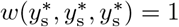. Visual inspection of the expression in (50) suffices to conclude that, for any value of the resident *θ*_s_, the most threatening value *τ*_s_ of any mutant type *τ* = (*τ*_d_, *τ*_s_) against the resident is the one that maximizes *W*_s_(*τ*_s_, *θ*_s_), i.e., such that the equilibrium strategy used in a mutant-mutant interaction, 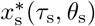, solves 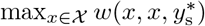. Given differentiability of *w* and *χ* = R, the strategy 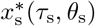 of the most threatening mutant must thus satisfy the following first-order condition:

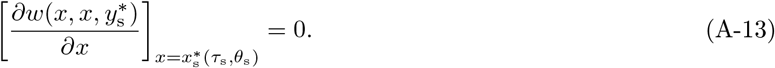

Since from eq. (46), 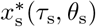 must satisfy the following first-order condition for utility maximization by a mutant,

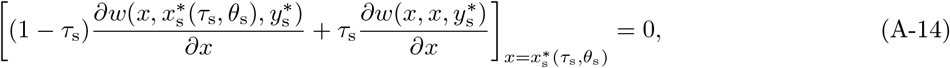

our assumption that 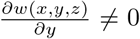 for all (*x, y, z*) ∈ *χ* ^3^ implies that *τ*_s_ = 1 is the unique value of *τ*_s_ ∈ [0, 1] which induces mutants to use the strategy that maximizes 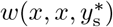. Hence, we can conclude that for any resident type *θ*, the most threatening mutant type has *τ*_s_ = 1. Note further that the argument developed above implies that *θ*_s_ = 1 implies *W*_s_(*τ*_s_, *θ*_s_) = 1 if *τ*_s_ = 1 and *W*_s_(*τ*_s_, *θ*_s_) <1 if *τ*_s_ <1. Hence, *θ*_s_ = 1 is sufficient for *W*_s_(*τ*_s_, *θ*_s_) ≤ 1 for any *τ*_s_ ∈ [0, 1].

We turn now to the term in eq. (49), which measures the individual fitness of a mutant in an interaction with a resident. Suppose that the resident type is 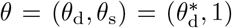, where 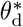 is defined in the statement of the result. By definition of 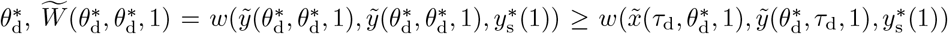 for all 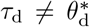. Since 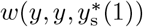 is strictly concave in *y* (owing to our assumption that the utility function is strictly concave in its first argument), which achieves its maximum for *y* satisfying (A-13), it follows that the value of *κ* ∈ [0, 1] that would maximize the function 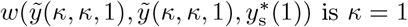.Together with the preceding inequality, we can thus conclude that 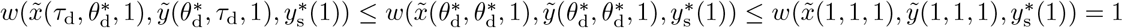. In other words, for any 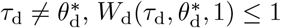.

The preceding conclusions, namely that *W*_s_(*τ*_s_, 1) ≤ 1 and 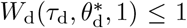, imply that whatever is the value of relatedness *r*(*τ, θ*), if the resident type is 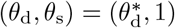, then *W*(*τ, θ*) ≤ 1 for any *τ* ≠ *θ*. **Q.E.D**.

## Footnotes

1 While we allow for individuals surviving from one demographic time point to the next, the survival probability is assumed independent of age, so that there is no effective age structure in the population.

2 When invasion fitness is differentiable, the quantities *S*(*θ*), *H*(*θ*), and *J* (*θ*) in fact allow for a complete classification of the singularities of the evolutionary dynamics (Geritz et al., 1998). Thus, when *H*(*θ*\*) *>* 0 and *J* (*θ*\*) <0, a singular type *θ*^*\**^is an evolutionary branching point; namely, an attractor of the evolutionary dynamics that subsequently splits the population into distinct morphs leading to the coexistence of different types in a protected polymorphism. When *H*(*θ*\*) <0 and *J* (*θ*\*) *>* 0 we have a so-called garden of eden state of the evolutionary dynamics, an uninvadable trait value that is unattainable by gradual evolution. Finally, if *H*(*θ*\*) *>* 0 and *J* (*θ*\*) *>* 0 then the singular type *θ*^*\**^is a an uninvadable repellor.

3 This follows from the fact that under the full evolutionary dynamic process of quantitative traits, the selection gradient *S*(*θ*) describes the direction of selection on small trait deviations regardless of population genetic states and demographic structures (Rousset and Billiard, 2000, Rousset, 2004, p. 206, Priklopil and Lehmann, 2021). This entails that any mutant invading the population when rare will eventually substitute the resident and recurrent mutations will drive the trait towards the singularity within its neighborhood when condition (3) is satisfied. This “invasion implies substitution” result was first noted in a special case by Hamilton (1964a) and called “a gift from god” (Hamilton, 1988). See also Eshel et al. (1997) for a different line of argument reaching the same conclusions.

4 Invasion fitness eq. (2) must be defined for all *τ* ∈ Θ including *τ* = *θ*. In such a monomorphic population, all individuals use strategy 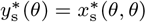 and thus the demographic consistency relation *W* (*θ, θ*) = 1 will be verified since 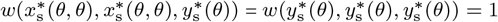 for all *θ* ∈ Θ.

5 An individual’s utility function is indeed simply a representation of its preferences. For any pair of strategies *x* and *y*, an individual’s preferences over available strategies tell whether the individual prefers *x, y*, or is indifferent between the two. Under certain conditions, such a preference ordering can be fully described by a function that associates a real number to each strategy, namely the utility function (see, e.g., Mas-Colell et al., 1995; Binmore, 2011). An individual is assumed to choose a strategy with the highest possible value of the function, since this is the strategy it prefers. Utility maximization is not to be taken literally: it is simply a mathematical tool used to describe behavior that amounts to choosing the preferred item from the strategy set. A pair of strategies then constitutes a Nash equilibrium if each individual uses a strategy which, given the other individual’s strategy, is the one it prefers.

6 This entails no loss of generality, and simply depends on how one defines the strategy set. For example, if the interaction at hand is a public goods game, then let *x* and *y* denote the contributions to the public good. If the interaction at hand is a common pool resource game, then let *x* and *y* denote the inverse of contributions to extracting resources.

7 Note that our model is different from the one in Alger and Weibull (2013), where a resident faces a positive probability of being matched with a mutant. In our model, the mutant trait appears initially in one single individual, and uninvadability obtains if the lineage created by this initial mutant goes extinct in the population (while invadability obtains if the lineage size created by this initial mutant becomes infinite and thus reaches positive frequency in the infinitely large population). During the time the mutant lineage is around in the population there can thus only be a finite number of mutants, and hence residents face a zero probability of being matched with a mutant in this infinitely large population. See Box 1 or Alger et al. (2020) for a formal explanation.

8 This may not be immediately apparent, for the results in Alger et al. (2020) (see Propositions 1 and 2) state as a necessary and sufficient condition for a utility function to be uninvadable, that the equilibrium strategy in a population where all individuals have this utility function be an uninvadable strategy (i.e., uninvadable in a setting where the set of traits is the set of strategies, a setting that Alger et al. (2020) call strategy evolution). One can check that the condition for *θ*\* = *r*(*θ*\*, *θ*\*) to be uninvadable in our setting, i.e., *H*(*θ*\*) <0 (see eq. (24)), coincides with the condition for the equilibrium strategy *y*\* (*θ*\*) to be an uninvadable strategy under strategy evolution. This is so because owing to eq. (12), 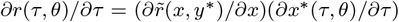, whereby eq. (24) becomes 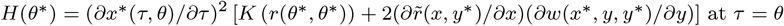; and under strategy evolution the invasion fitness of mutant type *x* in a monomorphic resident population *y* is 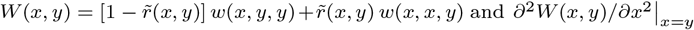 corresponds to the term in square brackets in *H*(*θ*\*).

9 Such independence was originally assumed in evolutionary game theory models with assortative interactions (Hines and May-nard Smith, 1978; Maynard Smith, 1982) and later used in preference evolution models (Alger and Weibull, 2013).

10 Because the behavioral perturbations of other-regarding utility functions, and thus *ρ*(*θ*), typically differ from the ones with the partially Kantian utility function, a singular *α* will typically differ from the singular Kantian coefficient, however.

11 Note that the definition of the function 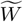 allows us to evaluate *W*_*d*_ under the hypothesis that the resident would apply the Kantian coefficient *θ*_d_ even when interacting with another individual with the same type. This is useful for the proof of the result, and for the argument developed in the next paragraph.

12 To see that polymorphism is likely under preference evolution consider Proposition 1 of Heifetz et al. (2007a), which establishes conditions for the evolutionary viability of pessimism or optimism for a particular individual fitness function. Using our notation, these preferences entail the utility function 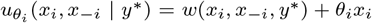, where *θ*_*i*_ is the evolving quantitative trait that can be taken to describe optimism when *θ*_*i*_ *>* 0 and pessimism when *θ*_*i*_ <0. It is straightforward to check that the singularity in Proposition 1 of Heifetz et al. (2007a) is both convergence stable and uninvadable, as it should under the measure dynamics they consider (e.g.,Cressman and Hofbauer, 2005). However, it is also straightforward to find parameter values of the fitness function they use Heifetz et al., 2007a, eq. 8 such that the singularity is convergence stable and invadable, and thus conducive to an adaptive polymorphism in dispositions, e.g., the coexistence of optimists and pessimists.

